# Quantifying the Tissue-Specific Regulatory Information within Enhancer DNA Sequences

**DOI:** 10.1101/2021.05.02.442309

**Authors:** Philipp Benner, Martin Vingron

## Abstract

Recent efforts to measure epigenetic marks across a wide variety of different cell types and tissues provide insights into the cell type-specific regulatory landscape. We use this data to study if there exists a correlate of epigenetic signals in the DNA sequence of enhancers and explore with computational methods to what degree such sequence patterns can be used to predict cell type-specific regulatory activity. By constructing classifiers that predict in which tissues enhancers are active, we are able to identify sequence features that might be recognized by the cell in order to regulate gene expression. While classification performances vary greatly between tissues, we show examples where our classifiers correctly predict tissue specific regulation from sequence alone. We also show that many of the informative patterns indeed harbor transcription factor footprints.

## 1 Introduction

Complex multicellular organisms comprise a large number of different cell types, which all share the same genome. Nevertheless, cell morphology and function are determined by the combination of genes that are expressed [Alberts, 2017, Ralston and Shaw, 2008]. To unravel how cells control gene expression, we first must identify all regulatory elements of the genome. This is relatively easy for promoters, because they are located proximal to the target gene’s transcription start site. Enhancers, which regulate cell type-specific gene transcription, are located distal to transcription start sites and therefore much more difficult to identify. Recent efforts focused on measuring epigenetic marks across a variety of different cell types and tissues [Stamatoyannopoulos et al., 2012, ENCODE Consortium, 2012, Bernstein et al., 2010]. These marks, including histone modifications, DNA methylation, and DNA accessibility, allow one to identify for each cell type regions of the genome that may act as enhancer elements.

However, this information alone can only be regarded as a first step towards understanding the regulatory program of a cell. Ultimately, we would like to understand why certain regions act as enhancer elements in particular cell types and how they drive gene expression. We believe that this information must be encoded in the genome in the form of binding sites for proteins such as transcription or pioneer factors. Hence, it should be possible to use the genomic DNA sequence not only for identifying enhancer elements but also for predicting the cell types in which the elements are active. Unfortunately, our knowledge of transcription factors including their DNA binding preferences and interactions remains incomplete. Instead, our goal is to quantify how much information about the cell type-specific activity of an enhancer element is actually encoded in the DNA sequence. For this, we may use enhancers identified from epigenetic marks and train classifiers that predict for each element the cell type in which it was found to be active. The classification performance then allows us to quantify to what extent there exists a correlate of cell type-specific epigenetic marks in the DNA sequence of enhancer elements.

An increasing number of studies focus on the prediction of regulatory elements from DNA sequence. Especially the DNA sequences of promoters have been studied in depth and several cell type-specific binding patterns have been identified [Roider et al., 2009, Halperin et al., 2009, Calo and Wysocka, 2013, Conlon et al., 2003, GuhaThakurta, 2006]. Since regulatory regions can be better targeted by transcription factors when they are not concealed by histones [Thurman et al., 2012], other studies focused on the prediction of accessible regions as measured by DNase-seq [Boyle et al., 2008] or ATAC-seq [Buenrostro et al., 2015]. Within cell types, accessible regions can be predicted from DNA sequence alone with high accuracy [Hashimoto et al., 2016]. Especially for ubiquitous regions, which are open in many or all cell types, the accuracy is very high. An analysis of feature importance revealed motifs of pioneer factors as well as CpG dinucleotide content [Hashimoto et al., 2016]. Other studies focused on the genome-wide prediction of active enhancers from DNA sequence [Lee et al., 2011, Yang et al., 2017], which were identified either through enhancer-specific patterns of histone marks or ChIP-seq experiments targeting EP300. The overall performance of such methods is good and comparable to the performance of open chromatin predictions. The focus of these studies, however, lies on the genome-wide prediction of enhancer elements [Lee et al., 2011, Hashimoto et al., 2016, Kelley et al., 2016, Yang et al., 2017] within a particular cell type. Methods developed for this task are trained to separate regulatory elements from other genomic sequences and may identify sequence patterns common to all regulatory elements or the genomic background.

Instead, we pursue a different line of research that focuses on learning tissue or cell type specific patterns [Kelley et al., 2016, 2018]. Our learning set-up consists of enhancer DNA sequences active in one or more tissues (positive set) versus enhancer DNA sequences from all remaining tissues (negative set). We use the recently developed leapfrog logistic regression Benner [2021] on sequence k-mers that allows to efficiently compute sparse solutions on very high-dimensional data. From the trained classifiers we quantify how much cell type-specific regulatory information is actually contained in the DNA sequence of enhancers. Furthermore, the classifiers are easy to interpret and allow to identify DNA subsequences or code words that may drive cell type-specific gene expression. We use ATAC-seq data to validate our findings by showing that many of the informative patterns indeed harbor transcription factor footprints (Figure 1).

**Figure 1.**
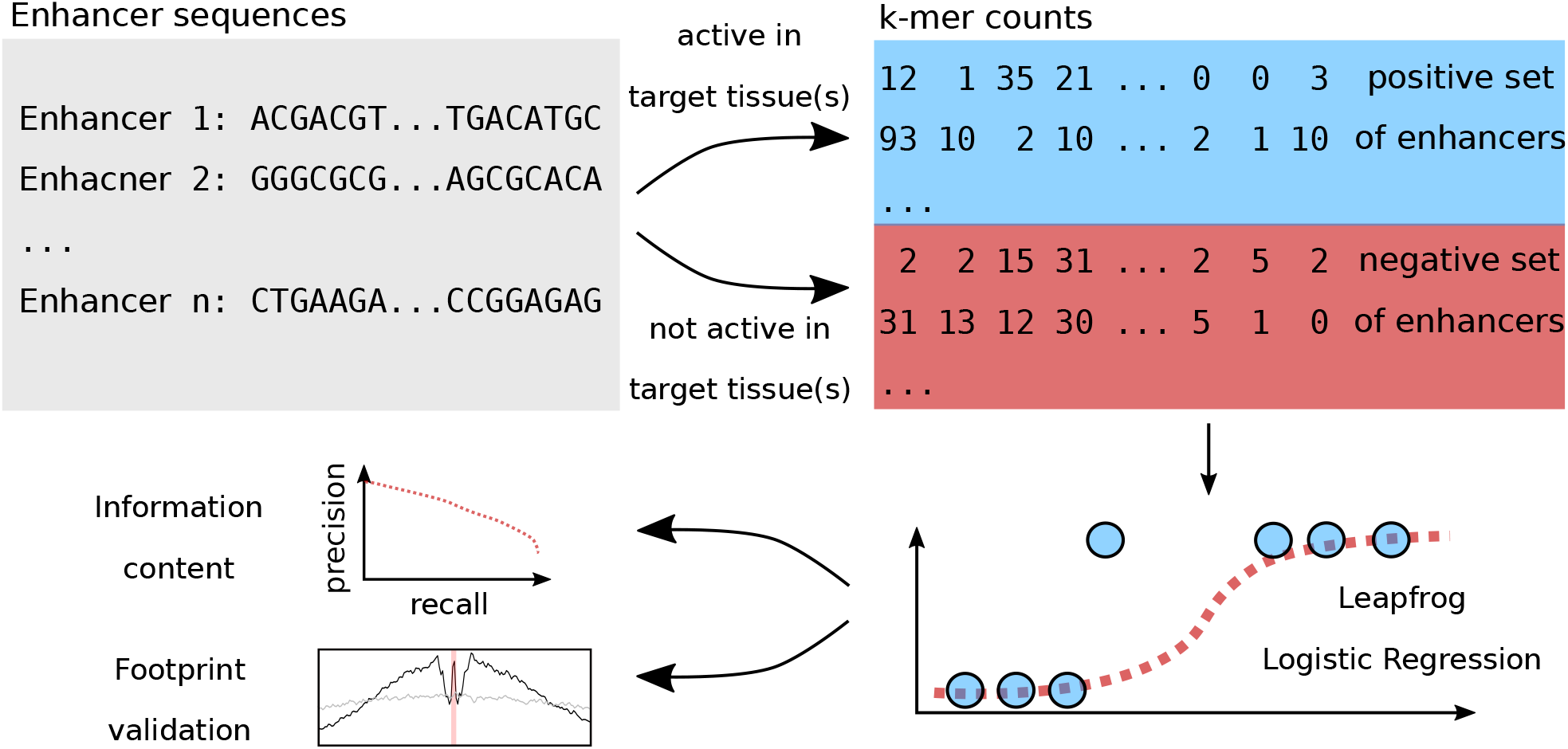
Overview. We use enhancer sequences from multiple tissues and extract k-mer occurrences. Logistic regression classifiers are trained with leapfrog regularization. From the performance of trained classifiers we quantify cell type-specific regulatory information within DNA sequences. Futhermore, we extract cell type-specific patterns and validate them through ATAC-seq footprints

Identified code words are only meaningful if the training set is of high quality and reflects our research objectives. Our interest lies in tissues as opposed to isolated cell lines that sometimes are derived from immortalized cancer cells. ENCODE provides comparable epigenetic data across several tissues in the mouse embryo. We use ModHMM [Benner and Vingron, 2020] for computing genome segmentations based on ATAC-seq, RNA-seq, and ChIP-seq data for multiple histone modifications. ModHMM computes highly accurate annotations of active enhancers that are not tainted by promoter elements or inactive (e.g. primed) enhancers.

## 2 Results

### 2.1 Characteristics of data sets

We obtained data from eight tissues (heart, kidney, liver, limb, lung, forebrain, midbrain, hindbrain) of embryonic mouse at day 15.5 from the ENCODE project (Supplementary Tables 26 and 27). ModHMM [Benner and Vingron, 2020] was used to compute genome segmentations for each of the tissues. The segmentations provide, among others, the genomic coordinates of several types of regulatory elements within each tissue, such as active promoters and enhancers as well as primed regions. Across tissues, regulatory elements that overlap are merged and we record the set of tissues in which the element was observed. To avoid biases, we set the length of all regulatory regions to 1000 bps around the center. Figure 2 shows the tissue specificity of promoters and enhancers. Most promoters are active in all 8 tissues, which supports similar findings from RNA-seq studies [Ramsköld et al., 2009]. We also find that the observed-to-expected CpG ratio [Gardiner-Garden and Frommer, 1987], referred to as CpG ratio in the following, grows steadily with the number of tissues in which promoters are active. On the other hand, most enhancers are active only in very few tissues and there is almost no enhancer active in all eight tissues. These results suggest that for genes that are active in multiple tissues there exists a distinct set of enhancers in each tissue that drives expression. Many enhancers and promoters in forebrain, midbrain and hindbrain are active in two other tissues (i.e. Figure 2 shows a peak for brain tissues at three), reflecting the fact that brain tissues are very similar and share many regulatory elements. The tissue specificity of primed enhancers is very similar to active enhancers (Supplementary Figure 3).

**Figure 2.**
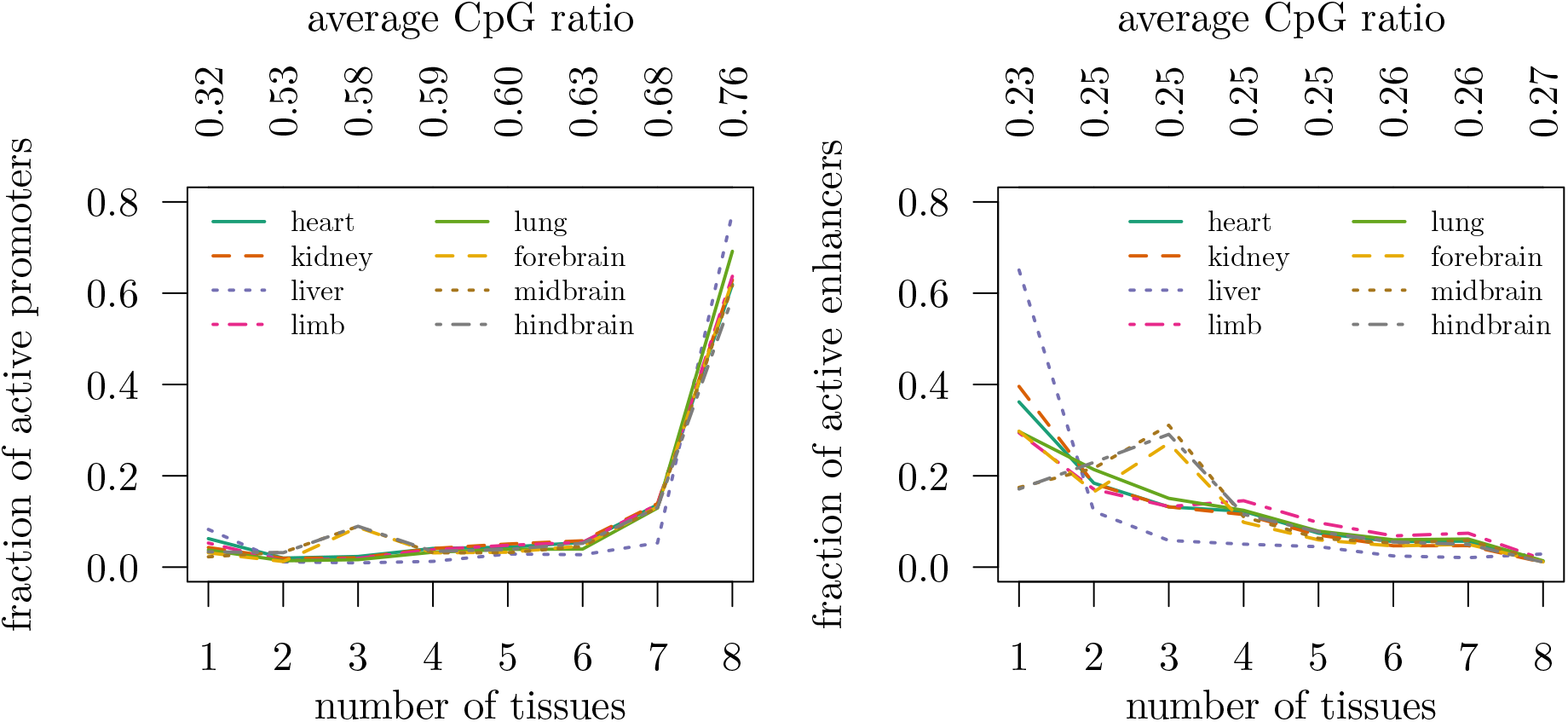
Tissue specificity of regulatory regions. The figures show the fraction of promoters (left) and enhancers (right) that are active in *n* ∈ {1, 2, …, 8} tissues, stratified by tissue. For instance, 80% of liver enhancers are active only in liver, but almost no enhancer is active in all eight tissues. In hindbrain around 30% of the enhancers are active in two other tissues. Above the figure the average observed-to-expected CpG ratio is shown

### 2.2 Learning set-up

As shown in the previous section, most enhancers are active in only a few tissues, which indicates that enhancers drive cell type-specific gene expression. A more detailed analysis on the location of the cell type-specific regulatory code can be found in Supplementary Section 1.3. Hence, we focus in the following on the analysis of enhancer DNA sequences.

For testing different feature sets, we constructed an unbalanced data collection using a one-vs.-rest strategy. It consists of eight data sets, each of which defines the enhancer elements of a single tissue as positive samples and the elements of all remaining tissues as negative. We refer to this collection as the *leaves data collection*. Furthermore, we constructed a second data collection where positive and negative sets consist of several tissues grouped together. The *clustered data collection* is defined in Table 1 and visualized as a tree in Figure 3a. For each data set in this collection, tissues were either assigned to the positive or negative set, based on a hierarchical clustering of enhancer regions (Section 4). An enhancer is considered a positive sample if it is active in any of the positive and none of the negative tissues.

**Table 1.** Definition of clustered data collection. Enhancer regions were clustered using a hierarchical clustering method. An edge of the resulting tree defines a bipartition of the tissues. Each data set in this collection corresponds to an inner edge of the tree

| data set | positive | negative |
| --- | --- | --- |
| 1 | liver | heart kidney limb lung forebrain midbrain hindbrain |
| 2 | kidney lung | heart limb liver forebrain midbrain hindbrain |
| 3 | heart kidney lung | limb liver forebrain midbrain hindbrain |
| 4 | forebrain midbrain hindbrain | heart kidney limb liver lung |
| 5 | midbrain hindbrain | heart kidney limb liver lung forebrain |

**Figure 3.**
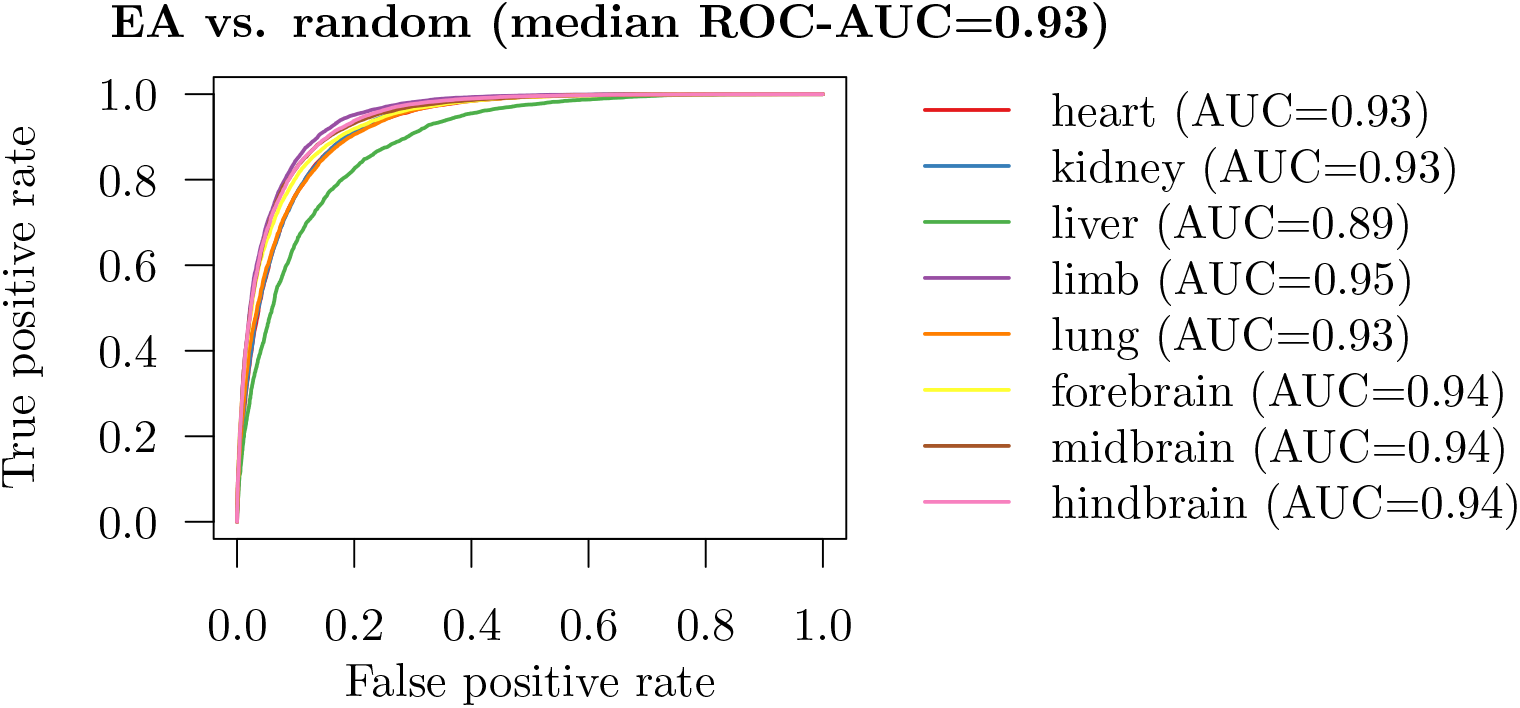
KLR classifier performance on active enhancers (EA) versus random regions. ROC curves were computed using 10-fold cross-validation

In this study, we mainly rely on logistic regression as classifier that we train with the recently developed leapfrog regularization (Section 4). We tested motif scores and k-mers as features, but found that k-mers generally yield better results (see Supplementary Section 1.1). For logistic regression with k-mers (KLR), we consider all k-mers of length 4-8 with any number of gaps (denoted by Ns). A k-mer and its reverse complement are considered equivalent, i.e. the feature vectors have a single coordinate for both, say, ANNTG and its reverse complement CANNT. We denote this pair as ANNTG|CANNT and refer to it as *code word*. The KLR classifier uses mainly code word counts as features. However, for extracting most important code words from our classifiers we only use code word occurrences as features, i.e. 1 if a code word appears in an enhancer sequence and 0 otherwise. This simplifies the interpretation of classifiers without much reducing their performance. To gauge the performance of the KLR classifier we use a support vector machine (SVM) with gapped k-mer string kernel [Lee, 2016].

### 2.3 Classifier choice and comparison

We first tested the performance of the KLR and SVM classifiers on active enhancers from the clustered collection. We tested the classifiers on each data set of the collection using 10-fold cross-validation (Supplementary Figure 4). Both classifiers perform relatively well with a median area under precision-recall curve (PR-AUC) of 0.71 and 0.67 respectively.

The performances of the KLR and SVM classifiers are similar and both could be used in this study. However, the KLR classifier is slightly better and much easier to interpret, i.e. the parameters of the KLR classifier can be directly linked to the importance of individual k-mers. The interpretation of SVMs, on the other hand, is much harder since there exists no natural way of extracting the importance of k-mers [Shrikumar et al., 2019]. In addition, training SVMs is computationally much more expensive. We therefore focus mainly on the KLR classifier in the following.

We then tested the KLR classifier on a positive set of active enhancers and a negative set of equal size consisting of random genomic regions. The random regions have the same length as the enhancers, i.e 1000 bps. We did not exclude any genomic regions, such as repetitive elements, for constructing the negative set. This ensures that our results are comparable to related studies [Hashimoto et al., 2016], which aim at whole genome predictions. The performance is overall very good across the eight different tissues with ROC-AUC values ranging from 0.90 to 0.97 and a median of 0.96 (Figure 3). This result is consistent with previous studies aiming at recognition of open chromatin regions [Lee et al., 2011, Hashimoto et al., 2016, Kelley et al., 2016, Yang et al., 2017]. Furthermore, we observed that, except for heart, important code words are highly AT-rich (Supplementary Tables 2 to 9).

To check the performance results we tested our cross-validation scheme on a control data set that contains enhancers from all tissues both as foreground and as background. The positive class consists of all enhancers on chromosomes 3, 5, 7, 11, 13, 17, and 19, the enhancers from the remaining chromosomes form the negative set. As expected, the classification performance on this data set is very close to random (Supplementary Figure 5).

### 2.4 Quantification of tissue-specific information

Our main interest is the quantification of tissue-specific information within enhancers. The construction of data sets is essential for extracting this information. If, for instance, we use enhancers active in a particular tissue as positive set and random genomic regions as negative set (as above), then it suffices that classifiers learn patterns common to all enhancers or the genomic background, because it is highly unlikely that the negative set contains enhancers that are active in other tissues. Most related studies [Lee et al., 2011, Hashimoto et al., 2016, Kelley et al., 2016, Yang et al., 2017] use random genomic regions as negative set. Instead, we train classifiers on data sets that contain DNA sequences of enhancers active in a particular subset of tissue (positive set) and inactive in all other tissues (negative set). This choice of the negative set is essential for truly learning tissue-specific information. Furthermore, we drop all enhancers that are active in both the positive and negative set, as those per se do not contain any information specific to tissues in the positive or negative set.

The classification performances can be interpreted as a lower bound on the tissue-specific information contained in the DNA sequences of enhancers. For quantifying this information we use the KLR classifier, a simple logistic regression on k-mers, because it performs as well as SVMs and at the same time is easy to interpret. We consider two different scenarios, either the positive set consists of multiple tissues (clustered data collection), or only a single tissue (leaves data collection).

The classification performance of the KLR classifier on the clustered data collection of active enhancers is shown in Figure 4. In general, the performances are much lower than for discriminating enhancers from random genomic regions. Separating brain from non-brain enhancers seems to be work best. Much harder is the task of separating hindbrain and midbrain enhancers from other tissues, mainly because hindbrain and midbrain enhancers seem to share many sequence features with forebrain enhancers.

**Figure 4.**
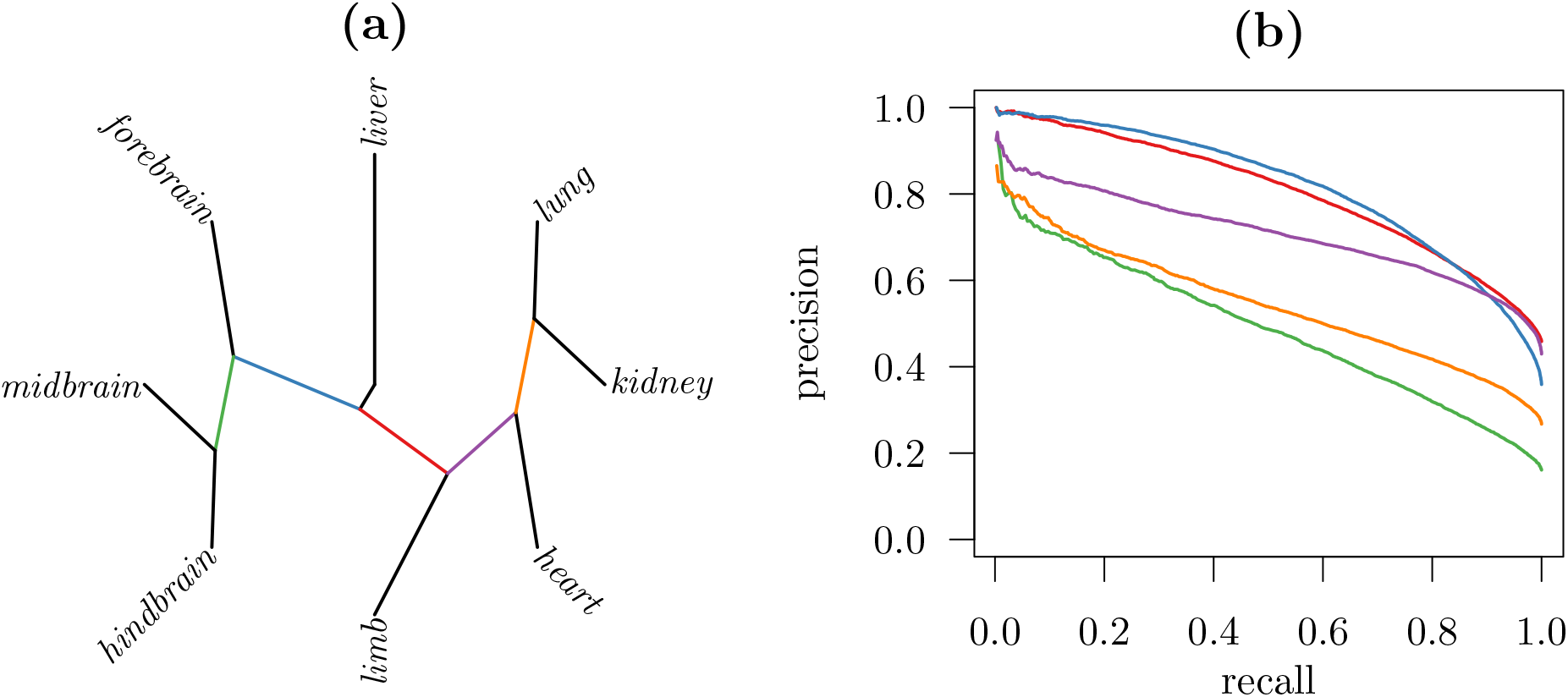
KLR classification performance on active enhancers from the clustered data collection. The left figure (a) shows a clustering of the 8 tissues. Every edge of the tree bipartitions the data into positive and negative samples. Respective classification performances using 10-fold cross-validation are shown on the right (b)

Even more difficult is the discrimination of active enhancers from a single tissue versus all other tissues. Figure 5 shows the performance of the KLR classifier on the leaves data collection. The prediction works best for forebrain and liver. The classifier trained on hindbrain shows the worst performance. This is surprising, because forebrain and hinbrain are both brain tissues, yet the classification performances are quite different. Nevertheless, we show in the following that all classifiers successfully identified important sequence patterns.

**Figure 5.**
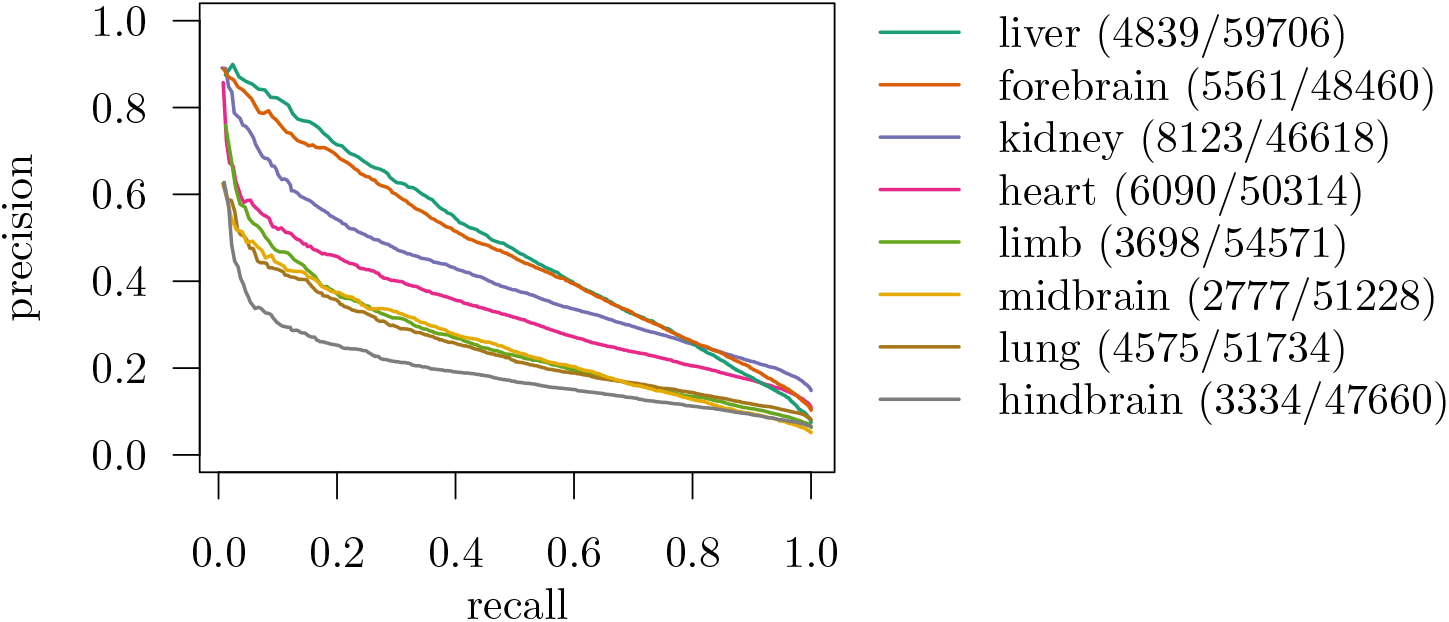
KLR classification performance on active enhancers from the leaves data collection. 10-fold cross-validation is used to evaluate precision-recall curves. The number of enhancers in the positive and negative (pos./neg.) set is reported in the legend

### 2.5 Code word counts versus occurrences

The KLR classifier can use as features either code word counts or occurrences. The former gives the number of times a code word is observed in the DNA sequence, while the latter is 1 if the code word is present and 0 if it is not. We wanted to understand if there is additional information in the code word counts, that is, if transcription factors recognize single binding sites or if the overall sequence affinity [Roider et al., 2006, Manke et al., 2008] is of importance. On the leaves data collection we found that the KLR classifier did not perform significantly worse if we only consider code word occurrences instead of counts (Figure 6). However, it is possible that the loss of count information is counteracted by the use of more code words. Indeed, we observe an increase in the number of features used by the optimal classifiers for enhancers (Supplementary Figure 8). For our purposes, using only code word occurrences is highly desirable, because it simplifies the interpretation of estimated classifiers, as discussed in the following section.

**Figure 6.**
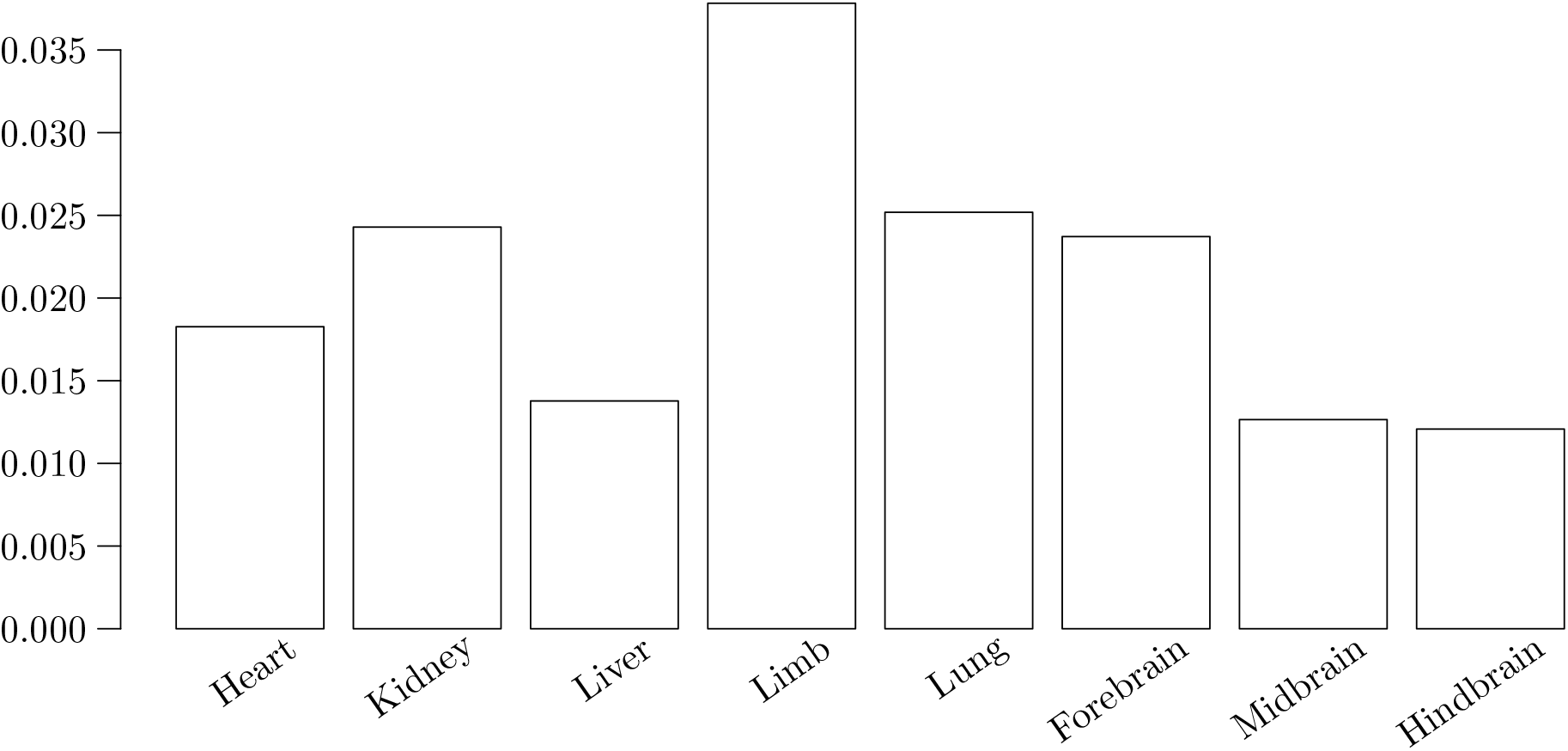
Code word counts versus occurrences. The figure shows the difference in area under precision recall (PR) curve when code word occurrences are used as opposed to code word counts. PR curves are evaluated for the KLR classifier using 10-fold cross-validation

### 2.6 Identification of cell type-specific regulatory code words in enhancers

To identify code words that might be recognized by transcription factors to drive cell type-specific gene expression, we use the KLR classifier applied to the leaves data set. The coefficients of the logistic regression can be understood as the importance of the corresponding code words. However, even when data is standardized, the interpretation of coefficients is much simpler when only code word occurrences are used [Gelman, 2008]. Therefore, we extract features from KLR classifiers that use only code word occurrences. Binarizing the data does not seem to lead to a significant drop in performance.

Features are extracted from classifiers trained on the leaves data collection, because those contain information about single tissues. We generally use 10-fold cross-validation to evaluate predictive performances. Features are extracted by first merging the 10 classifiers estimated during cross-validation. For each feature we take the coefficient with the minimum absolute value across all classifiers, which has the effect that only those features remain that have a non-zero coefficient across all training sets. We report only code words with positive coefficients, because those correspond to k-mers relevant to a single tissue, i.e. the positive set.

Table 2 shows the first two most important code words, a complete list is provided in Supplementary Tables 10 to 17. We used Tomtom [Bailey et al., 2009] to search for matching motifs in the Jaspar database [Desjardins and Naya, 2016]. For kidney, we identified a code word that may belong to PPARG-g, ER1, or the RXR-a/VDR heterodimer, the latter has known functions in kidney [Sugawara et al., 1997]. The limb classifier identified a code word that probably belongs to NFI-C binding sites, which is highly expressed in skeletal muscle cells [Chaudhry et al., 1997]. For heart and liver the most important code words seem to belong to the GATA family of transcription factors, while for lung we identified a code word that is most likely recognized by members of the Fox family. The top forebrain code word is a known binding site for NeuroD1, a neurogenic differentiation factor [Gao et al., 2009]. We also identified code words that are not associated with any transcription factor. Some contain two or more gaps (e.g. TAATNAA|TTNATTA or ATNNATCA|TGATNNAT), which might reflect co-binding sites of transcription factors.

**Table 2.**
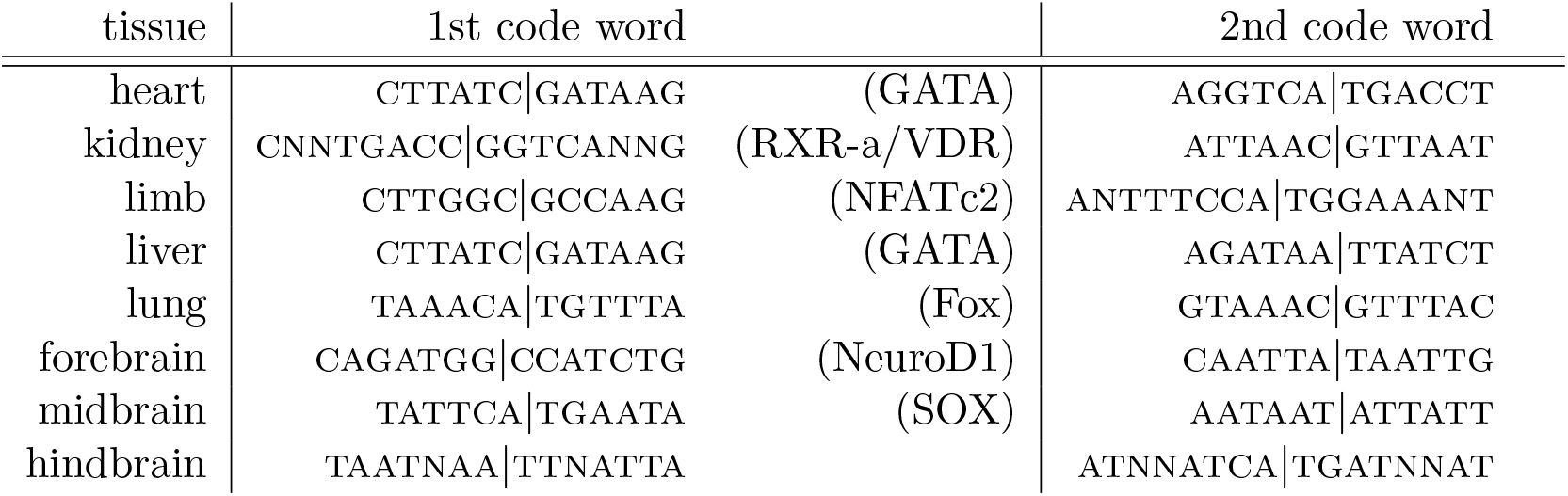
Tissue-specific code words. Two most important tissue-specific code words in active enhancers extracted from KLR classifiers with 100 non-zero coefficients (*q* = 100). Corresponding names of transcription factors are given in brackets if a unique assignment was possible

As a further control, we looked at ATAC-seq footprints [Li et al., 2019] around code words occurrences in active enhancers. We first computed the ATAC-seq coverages by treating paired-end reads as single-end and reducing read lengths to 2 bps. Forward strand reads where shifted by 4 bps, reverse strand reads by -5 bps. For each tissue, we scanned all active enhancer sequences for occurrences of the most important code word. Enhancers that do not contain the most important code word were omitted. We then aligned the ATAC-seq signal around the code word positions. When an enhancer contained the code word more than once, the position of the first occurrence was used. Except for forebrain, we observe clear ATAC-seq footprints at the centers (Figure 7), which suggests that the identified code words are indeed recognized by transcription factors. We used the same aligned sequences to compute logos of code word neighborhoods and observe almost no signal outside the code words (Supplementary Figure 10).

**Figure 7.**
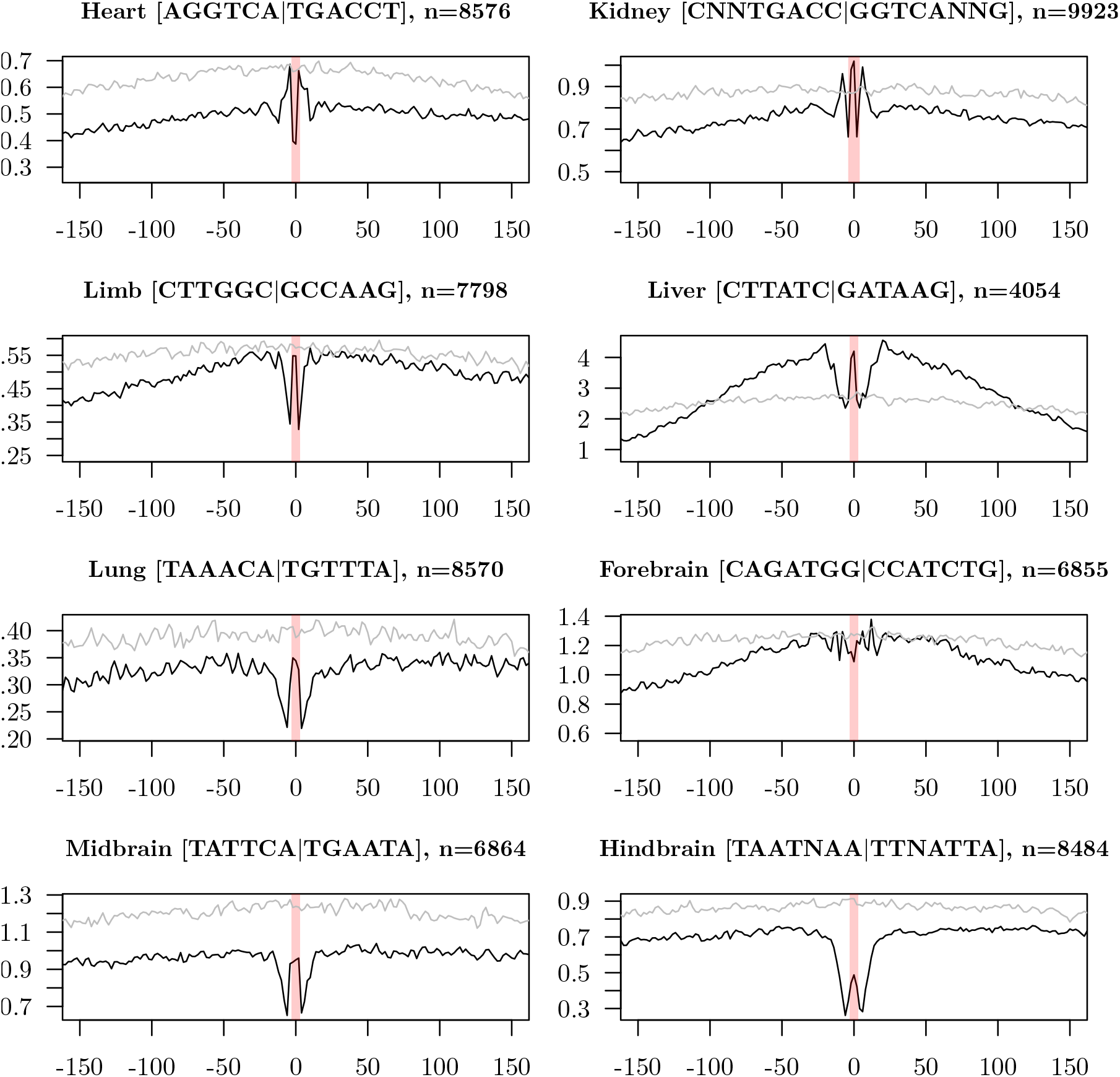
ATAC-seq footprints around most important code words of active enhancers (black lines). Regions are aligned around the center of code word occurrences and the footprint is computed as the average over all *n* active enhancers in the given tissue. The distance in bps from the center is shown on the x-axis. Red shaded areas show the positions of the code words. The control (gray lines) shows the average ATAC-seq signal of the same regions, but aligned at the center of the enhancer elements.

### 2.7 Sliding window predictions at selected loci

To further test our classifiers, we looked at several loci and computed sliding window predictions of active enhancers. More specifically, a sliding window of 2kb is used to compute predictions along the genome. We first use a classifier that discriminates between enhancer elements and random genomic regions. A second classifier separates brain from liver enhancers. All loci that we used for testing predictions were excluded from the training set of both classifiers. By combining the two classifiers we obtain predictions for active enhancers in brain along the genome. Figure 8 shows a region between Cdh9 and Cdh10, which in humans is known to harbor several enhancer elements and where genetic variants are associated with autism spectrum disorders [Wang et al., 2009, Inoue and Inoue, 2016]. The genome segmentation identified two enhancers that are active only in forebrain.

**Figure 8.**
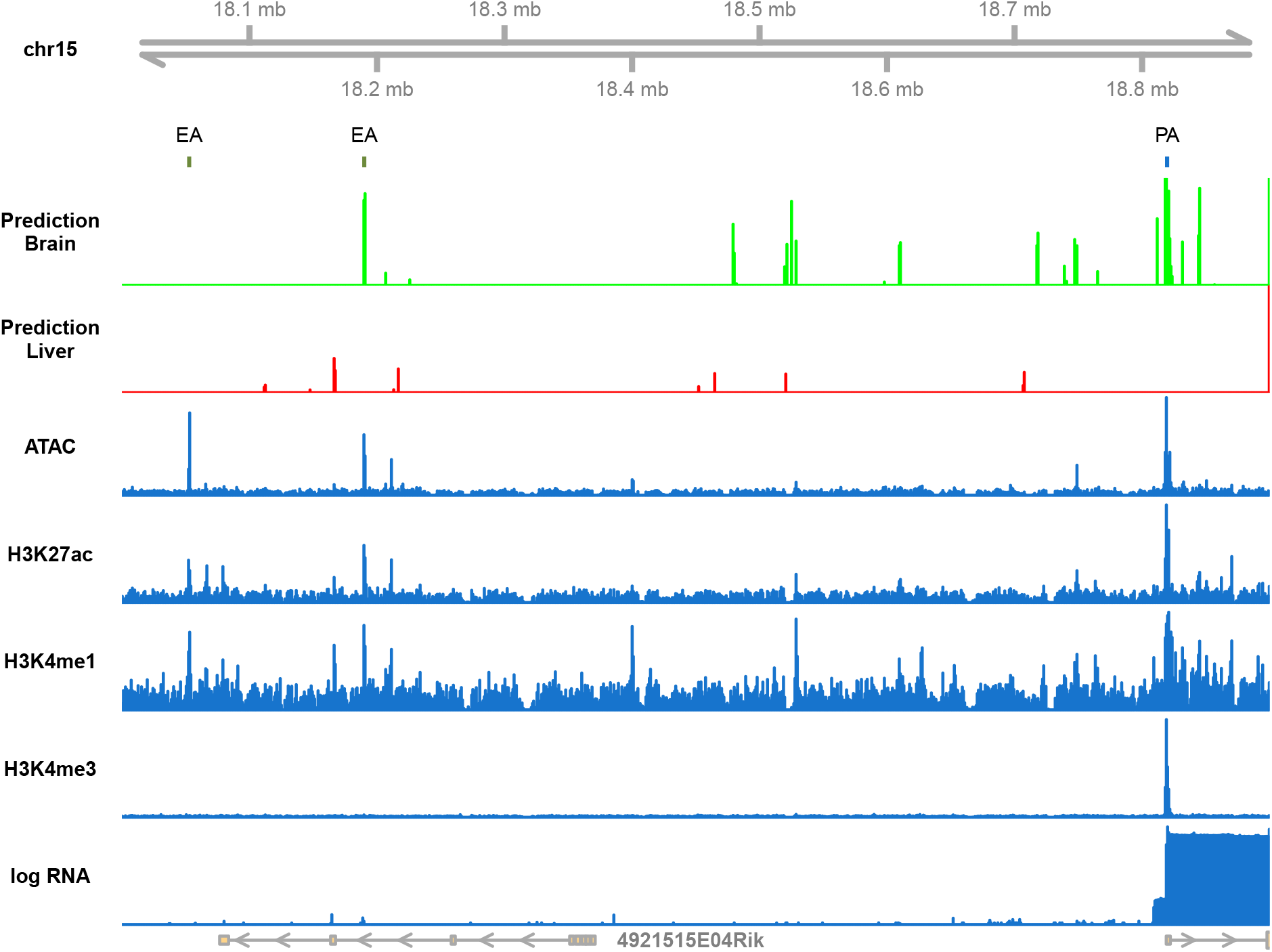
Sliding window predictions of active enhancers from DNA sequence. ModHMM-predicted active promoters (PA) and enhancers (EA) in forebrain are marked by small bars in blue and green respectively. Predictions of active brain enhancers from DNA sequence are shown in green, predictions of liver enhancers in red. The blue tracks below show ATAC-seq, histone marks, and RNA-seq coverages in forebrain.

The sliding window prediction of brain enhancers shows a peak at only one of the active enhancers, but also very few false positives within this locus. This observation is in line with our previous results. We showed that random genomic regions can be easily distinguished from enhancers, but differentiating between active enhancers in particular cell types is hard. However, the sliding window predictions also detect the active promoter of Cdh10, possibly because we did not train the classifier to discriminate between promoters and enhancers. Nevertheless, the precision of predictions is very promising and rarely reported in the literature. By inverting the predictions of the second classifier (probability of complement), we obtain enhancer predictions for liver. The prediction of liver enhancers shows no peaks at the brain enhancers or the Cdh10 promoter. Throughout this locus, the probability of liver enhancers is very low.

## 3 Discussion

In this study, we constructed classifiers that predict tissue-specific enhancer activity from DNA sequence. This has been done by several other studies [Lee et al., 2011, Hashimoto et al., 2016, Kelley et al., 2016, Yang et al., 2017], which focus on the genome-wide prediction of enhancers. We show that this task is relatively easy, because such classifiers mainly learn to discriminate between enhancer elements and other genomic regions. Instead, we focus on predicting cell type-specific activity of enhancer elements, which we achieve by training classifiers that discriminate between enhancers active in selected tissues. We use the classification performance as a proxy to measure how much information about cell type-specific activity is contained in the DNA sequence of enhancers. By using classifiers that are easy to interpret, we were able to extract important regulatory code words that might be recognized by transcription factors for driving cell type-specific gene expression. The ATAC-seq signature around identified code words show a clear pattern, which indicates that we indeed identified functional binding sites. Furthermore, the accuracy of our sliding window predictions is very promising and rarely reported in the literature.

The classification performance of active enhancers strongly depends on the tissues we want to discriminate. Our classifier performs well when discriminating between highly dissimilar tissues, such as brain and non-brain tissues. However, especially when the positive class consists of enhancers from a single tissue, the performance drops in many cases. One possible reason is that the tissues we are dealing with contain too many different cell types or that the data is simply too noisy. Another explanation is that we still have only a poor understanding of the cell type-specific regulatory code and the features required for predicting the activity of enhancer elements. It could be that not all the required information is contained in the DNA sequence of isolated enhancers and that we are still missing a piece of the puzzle.

There are many possible future research directions. For instance, single-cell ATAC-seq data could help to distinguish between cell types within a tissue and thereby help to train better classifiers. Furthermore, we know that transcription factors and other proteins bound to enhancers and promoters interact in order to initiate transcription. It is possible that the regulatory code is distributed among enhancers and promoters and that both must be considered jointly when training classifiers.

## 4 Methods

### 4.1 Enhancer identification with ModHMM

ModHMM [Benner and Vingron, 2020] is a hidden markov model for comuting genome segmentations similar to ChromHMM [Ernst and Kellis, 2012] and EpiCSeg [Mammana and Chung, 2015]. It uses a fixed number of hidden states and computes segmentations based on 8 features, namely ATAC/DNase-seq and RNA-seq in addition to histone modifications H3K4me1, H3K4me3, H3K27ac, H3K27me3 and H3K9me3. Compared to other genome segmentation methods, ModHMM better discriminates between different types of regulatory elements, such as active promoters and enhancers, by incorporating prior knowledge into the model. A reliable discrimination between enhancers and promoters is vital to this study.

ModHMM version 1.2.3 was used to compute genome segmentations of 8 embryonic mouse tissues with a bin size of 200 base pairs. All segmentations are available online at https://github.com/pbenner/modhmm-segmentations, from which we obtained the tissue-specific positions of active enhancers. For each type of regulatory element the 8 lists of tissue-specific genomic positions were merged by joining all elements that overlap by at least one bin. The size of all elements was then set to 1000 base pairs around the center and we acquired the DNA sequences of all regulatory elements from the mm10 assembly.

### 4.2 Logistic regression classifiers

We use leapfrog regularization [Benner, 2021] for estimating logistic regression classifiers on k-mers (KLR). The parameters *θ* ∈ ℝ^*m*+1^ of the classifier are estimated by maximizing the *ℓ*_1_-penalized log-likelihood function

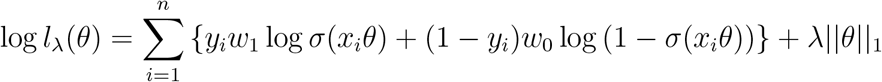

with class weights

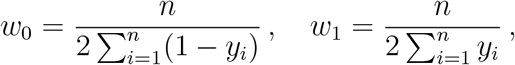

on a set of *n* observations *x* = {*x*_1_, *x*_2_, …, *x*_*n*_} with labels *y*_*i*_ ∈ {0, 1}. Each *x*_*i*_ = (1, *x*_*i*1_, *x*_*i*2_, …, *x*_*im*_) is a row vector of length *m* + 1, where *m* is the number of features. In the above formula, *σ* denotes the sigmoid function. *λ* ∈ ℝ_≥0_ is a parameter that controls the strength of the penalty and we use leapfrog regularization to determine its value so that 1*θ*1_0_ = *q* + 1 for some *q* ≤ *m*. Please note that although we wrote ||*θ*||_1_ in the above formula, we actually do not regularize the first component of *θ*, i.e. 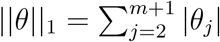. To compute maximum likelihood solutions, we implemented a just-in-time variant [Schmidt et al., 2017] of the SAGA algorithm [Defazio et al., 2014].

For the KLR classifier the set of features consists of all k-mers of length 4-8 with any number of gaps (denoted N). Reverse complements are considered equivalent, i.e. the vector *x*_*i*_ has a single coordinate for both ANNTG and its reverse complement CANNT, denoted ANNTG|CANNT. Hence, each *x*_*i*_ is a vector of dimension 156, 570 + 1 (*m* = 156, 570). We consider either code word counts or occurrences (binarized counts). In the case of code word counts, we first normalize the data to unit variance. The data is not centered at zero to retain the sparse structure.

For the entire study, classifier performance was evaluated using 10-fold cross-validation. We do not know the optimal number of features (non-zero coefficients). Therefore, we tested the performances for several values of *q*, which can be efficiently computed with leapfrog regularization. For the KLR classifier we always used *q* = 10, 100, 200, …, 900, 1000, 2000, …, 6000. 10% of the training data is used as a validation set for selecting the best *q*.

Our software is available at https://github.com/pbenner/kmerLr

### 4.3 K-mer SVM

LS-GKM [Lee, 2016] is a support vector machine (SVM) that also uses gapped k-mer frequencies as features. We mainly use it to benchmark the performance of the KLR classifier. However, the decisions that SVMs make are difficult to interpret and extracting important features from it is much more complicated than for the KLR classifier.

### 4.4 Clustering of tissues

The clustering of tissues is computed from ModHMM segmentations, where only the enhancer state is used. First, a similarity matrix between the tissues is computed by counting the number of overlapping enhancers between each pair of tissues. Afterwards, this matrix is used to compute a hierarchical clustering (based on the *hclust* method in *R*). The clustered data collection is computed from this clustering in the following way. By removing an inner edge from the hierarchical clustering tree, a bipartition of tissues is generated. This bipartition corresponds to a single data set of the collection and is used for labeling the enhancers as positive or negative. Enhancers that appear in both the positive and negative sets are removed.

## 5 Competing interests

The authors declare that they have no competing interests.

## 6 Acknowledgements

We thank Stefan Haas and Kirsten Kelleher for their comments on the manuscript and many inspiring discussions.

PB was supported by the German Ministry of Education and Research (BMBF, grant no. 01IS18037G).

## Notes

### Competing Interest Statement

The authors have declared no competing interest.

