## Supplementary Material for "Quantifying the Tissue-Specific Regulatory Information within Enhancer DNA Sequences"

### 1 Supplementary Figures and Tables

#### 1.1 SLR classifier

We tested a second type of logistic regression, called SLR in the following, which is based on motif scores. For the SLR classifier we use transcription factor binding motifs from the JASPAR 2018 database Khan et al. [2017], which contains $m = 119$ motifs for mus musculus. Given the DNA sequence of a candidate regulatory element, we compute the maximum motif score for each of the $m$ motifs. All scores are standardized before the coefficients $\theta$ are estimated.

We first tested the performance of the SLR classifier on active enhancers from the clustered collection (Supplementary Fig. 2). The SLR classifier with a median area under precision recall curve (PR-AUC) of 0.65 performs much worse than the KLR and SVM classifiers, with a median PR-AUC of 0.75 and 0.73 respectively. One reason for the poor performance of the SLR classifier could be that the set of motifs is incomplete. However, it may also be the case that motifs were estimated with inaccurate methods [Weirauch et al., 2013].

#### 1.2 Discriminating between active and primed enhancers

It has been observed from genome-wide analyses of histone marks that enhancers can be in several different states. For instance, primed enhancers are lacking histone mark H3K27ac and are thought to be activated later during development [Barski et al., 2007, Heintzman et al., 2007, 2009, Heinz et al., 2015, Calo and Wysocka, 2013]. We tested whether the KLR classifier is able to discriminate between primed and active enhancers within a tissue. Our results show similar classification performances for primed enhancers (Supplementary Fig. 6) as for active enhancers across all tissues. We extracted code words predictive for active enhancers (Supplementary Tables 18 to 25) and found that many are similar to known transcription factor motifs.

#### 1.3 Location of the cell type-specific regulatory code

To compare classification results across different types of regulatory elements (i.e. active CpG-low/CpG-high promoters and active enhancers), it is important to have an equal number of positive and negative samples of each type. We therefore constructed a *balanced data collection* (Table 1) consisting of three different data sets. We constructed the data collection so that each set has 800 positive and 800 negative samples for each type of regulatory element. The data collection contains only three data sets, because most CpG-high promoters are differential between brain and non-brain tissues. We use the CpG ratio with a threshold of 0.4 to separate CpG-high from CpG-low promoters.

| data set | positive | negative |
|---:|---:|---:|
| 1 | forebrain midbrain hindbrain | kidney lung |
| 2 | forebrain midbrain hindbrain | limb liver |
| 3 | forebrain liver | heart kidney limb lung |

Supplementary Table 1: **Definition of balanced data collection.**

Several different types of regulatory elements are known, each having a different role in the regulatory program. For instance, genes with high phyletic age tend to be expressed in many cell types and typically have promoters with high CpG dinucleotide content (CpG-high) [Zhu et al., 2008]. Their regulatory program is likely to be different from that of other promoters since the expression of target genes can be silenced by methylation of the CpG dinucleotides within the promoter [Bird, 2002]. On the other hand, genes that evolved more recently tend to be tissue-specific and their promoters show a low CpG dinucleotide content (CpG-low) [Zhu et al., 2008]. Additionally, much evidence exists to support the idea that cell type-specific expression is mainly regulated by enhancers [Voss et al., 1986, Heinz et al., 2015].

To verify these observations, we trained and tested the KLR classifier separately on CpG-high and CpG-low promoters as well as active enhancers from the balanced data collection. Comparing the performances of the three sets of classifiers allows us to speculate about how the regulatory code is distributed over the elements. Indeed, we observe that the classification performances vary strongly across the three types of regulatory elements (Fig. 1). The performance is highest for enhancers and CpG-high promoters. This result suggests that the regulatory code is mainly embedded in enhancers, which then drive the activity of CpG-low promoters. CpG-high promoters are known to be a special case, since they can be silenced through CpG methylation [Deaton and Bird, 2011]. Our results suggest that the regulatory code for CpG-high promoters is, at least to some extent, present in the promoter DNA sequences and not outsourced to enhancers. However, we also know that the distribution of the CpG ratio of active promoters varies greatly between tissues (Supplementary Figs. 6 and 7) [Roider et al., 2009], with brain tissues having the highest CpG ratios.

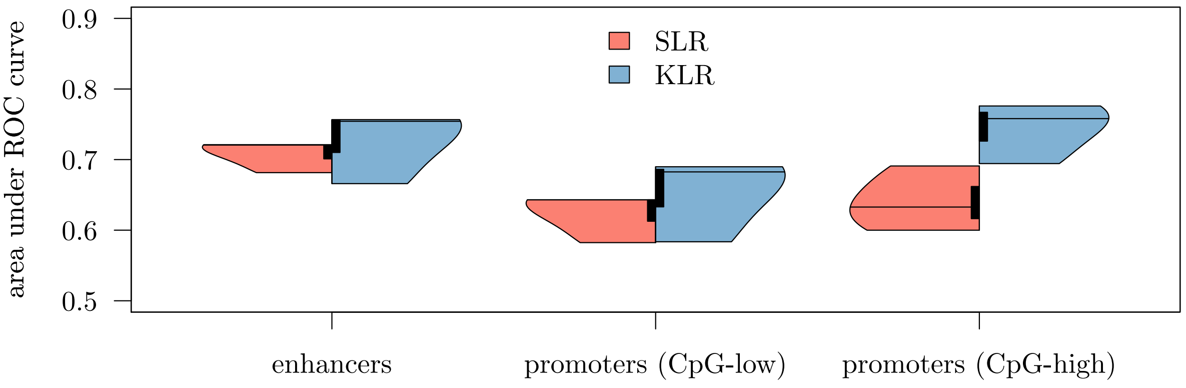


Supplementary Figure 1: **Classifier performances stratified by type of regulatory elements.** The figure shows the distribution of area under ROC-curves using 10-fold cross-validation for the SLR and KLR classifiers

#### 1.4 Importance of combinatorial binding of transcription factors

It is well established that in eukaryotes, interactions between transcription factors are essential for the regulation of gene expression, either by direct protein-protein interactions between factors or by interactions through the basal transcription machinery [Johnson and McKnight, 1989, Mitchell and Tjian, 1989, Wasylyk et al., 1990, Janson and Pettersson, 1990, Kel et al., 1995]. The combinatorics of transcription factor binding within regulatory elements have been studied extensively for promoters [Pilpel et al., 2001, Zhu et al., 2005] and enhancers [van Bömmel et al., 2018]. We are interested in determining to what extent combinatorial binding is important for predicting cell type-specific activity of regulatory elements. To investigate this question, we extended the SLR and KLR classifiers to include interaction terms.

Both the KLR and SLR classifiers can model co-occurrences by adding interaction terms to the logistic regression, i.e. we define

$$x \odot \theta = \theta_0 + x_{i1}\theta_1 + \cdots + x_{im}\theta_m + x_{i1}x_{i2}\theta_{m+1} + x_{i1}x_{i3}\theta_{m+1} + \cdots + x_{im-1}x_{im}\theta_{(m+1)m/2}\,,$$

where $\theta \in \mathbb{R}^{(m+1)m/2+1}$. For the KLR classifier, $m = 156,570$ so that $(m+1)m/2 = 12,257,160,735$, which is computationally too expensive to estimate. Instead, we first select the 1000 most important code words and consider co-occurrences only between those, resulting in a parameter vector $\theta$ of dimension $500500+1$. This procedure is equivalent to assuming *strong hierarchy*, which is not only computationally necessary, but often argued to be sensible [Bien et al., 2013].

We compared the classification performance of both classifiers with and without interaction terms on the leaves data collection. Our results show that there is no significant difference between the two types of classifiers (Fig. 2), although classifiers that model co-occurrences show mainly non-zero coefficients for interaction terms. For the SLR classifier we observe a very marginal increase in classification performance on enhancers when interactions are considered, but the opposite is the case for the KLR classifier. This result might suggest that combinatorial binding is not important for cell type-specific activity of regulatory elements. However, it may also be the case that transcription factors generally co-bind in close proximity so that this effect is already captured by the SLR and KLR classifiers without interaction terms.

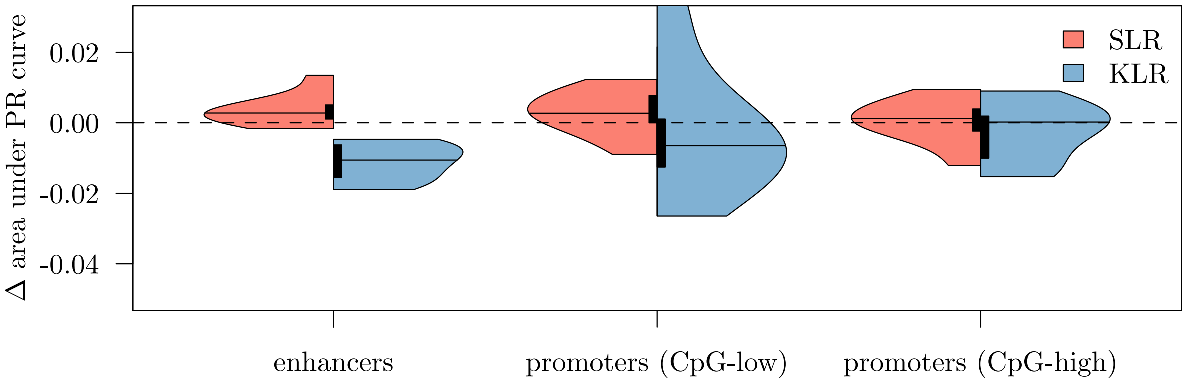


Supplementary Figure 2: **Importance of combinatorial binding.** Difference in area under precision recall (PR) curve between classifiers with and without interaction terms

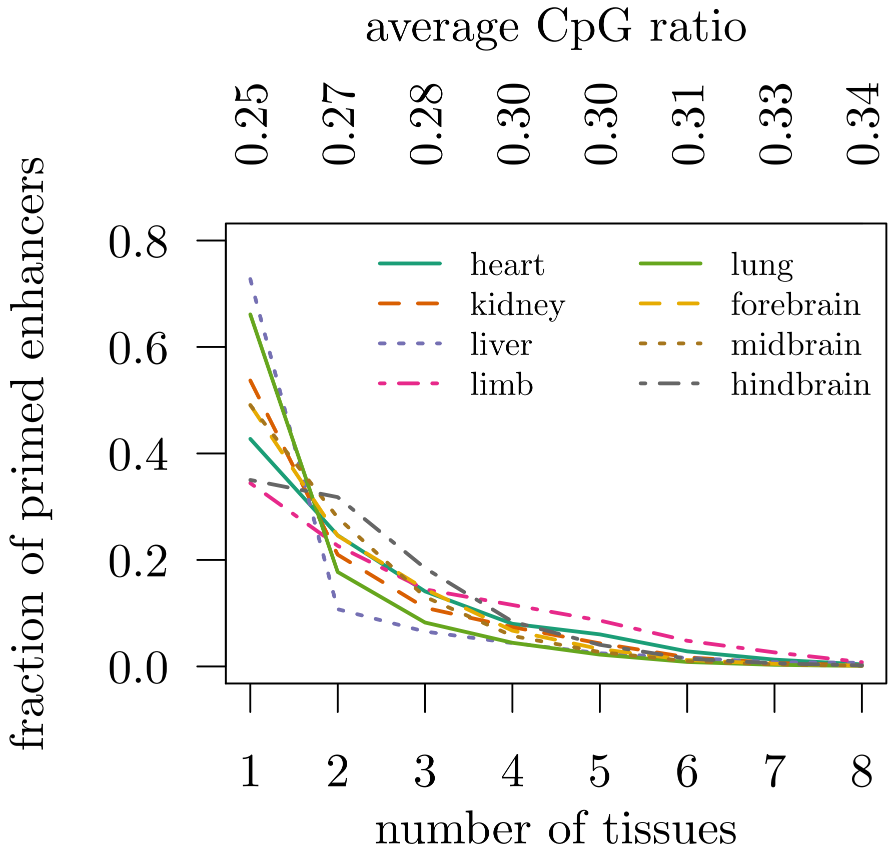


Supplementary Figure 3: Tissue specificity of primed enhancers

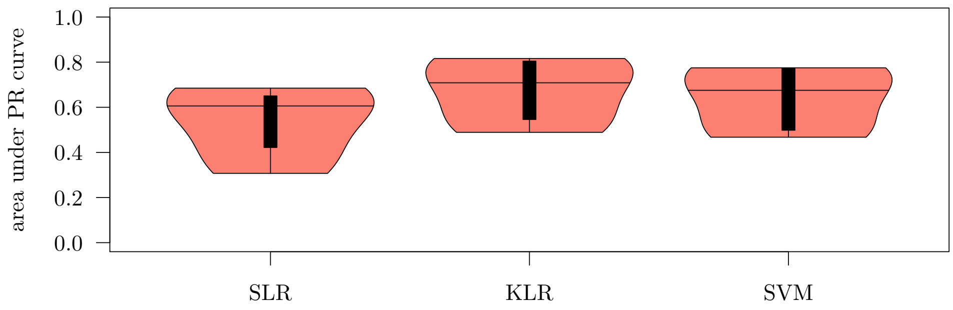


Supplementary Figure 4: Classifier performance evaluated on clusted data collection of active enhancers

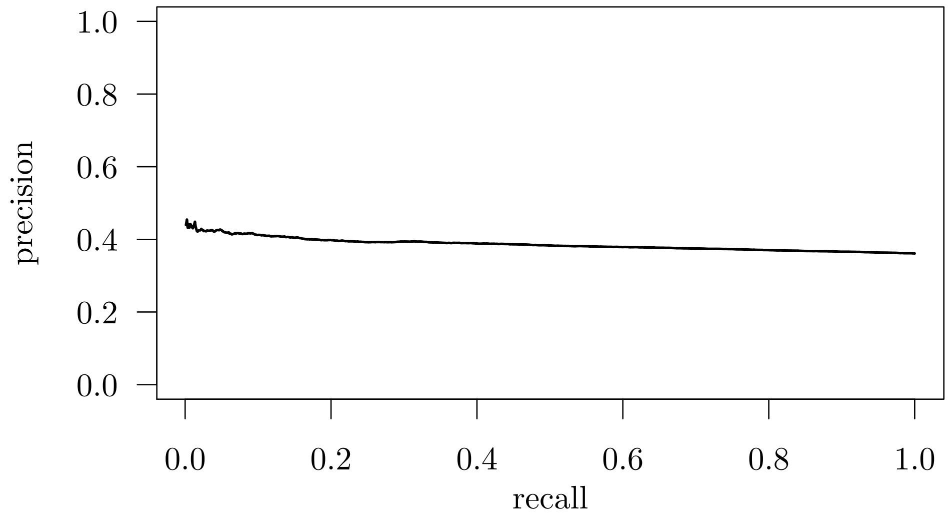


Supplementary Figure 5: KLR classifier performance on control data set

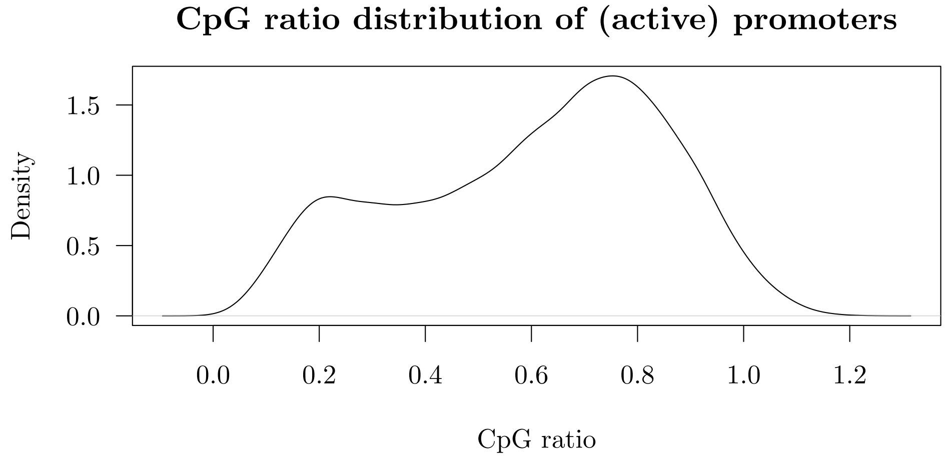


Supplementary Figure 6: CpG ratio distribution of promoters, active in at least one tissue

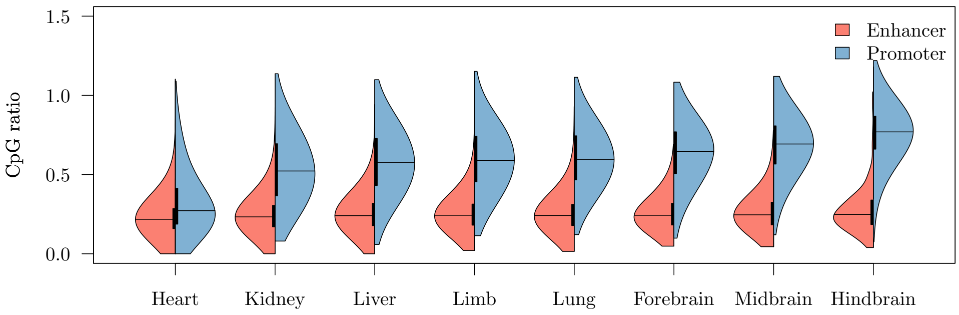


Supplementary Figure 7: CpG ratio of active enhancers and promoters stratified by tissue

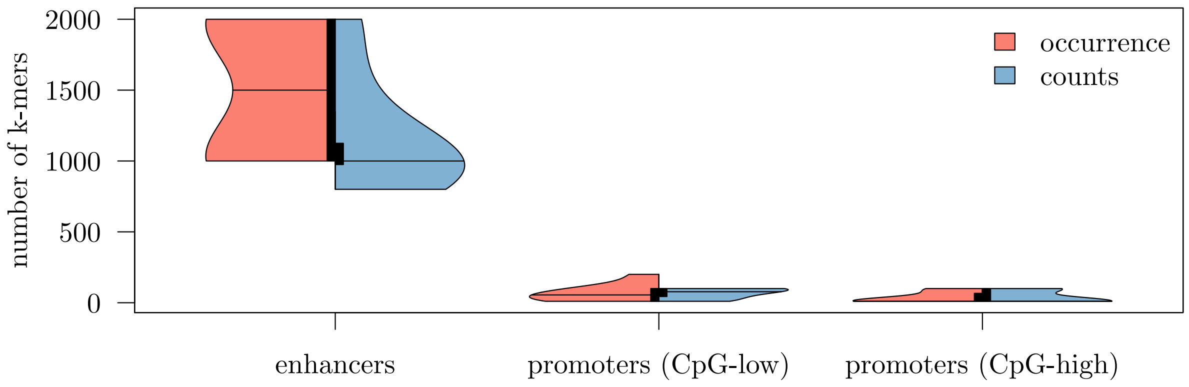


Supplementary Figure 8: Number of features used by the optimal KLR classifier when using code word occurrences or counts

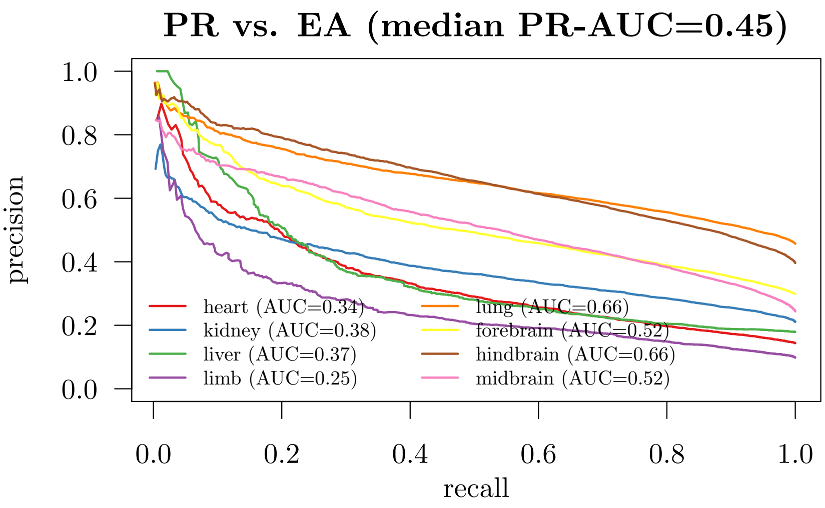


Supplementary Figure 9: KLR classification performance on primed (PR) versus active (EA) enhancers within tissues

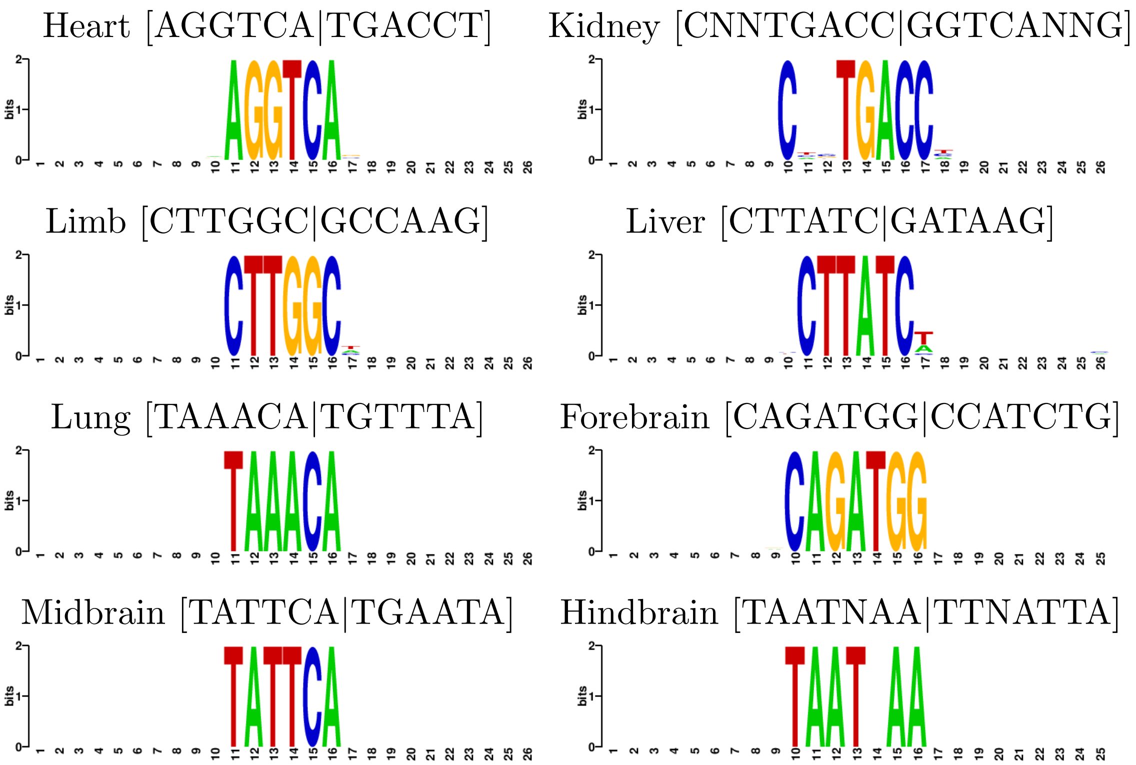


Supplementary Figure 10: Sequence logos of code word neighborhoods in active enhancers.

| Coefficient | k-mer |
|---|---|
| 1.220076e-01 | CNGCNCNG\|CNGNGCNG |
| 1.133865e-01 | CNGCNGNC\|GNCNGCNG |
| 1.104353e-01 | CCCNGCC\|GGCNGGG |
| 1.092400e-01 | GCNCNGC\|GCNGNGC |
| 9.447861e-02 | CNGNCAGC\|GCTGNCNG |
| 9.439475e-02 | CCGNNC\|GNNCGG |
| 8.421516e-02 | CNGCNGC\|GCNGCNG |
| 8.076685e-02 | CNCCNCCC\|GGGNGGNG |
| 8.025976e-02 | CGNNNGC\|GCNNNCG |
| 7.957235e-02 | CGCNG\|CNGCG |
| 7.945013e-02 | CAGNGCNG\|CNGCNCTG |
| 7.579287e-02 | CNCNCNGC\|GCNGNGNG |
| 7.021016e-02 | CCTNCCNG\|CNGGNAGG |
| 6.449539e-02 | CGNGC\|GCNCG |
| 5.339787e-02 | CNGGCTNC\|GNAGCCNG |
| 4.705916e-02 | CAGCNCNG\|CNGNGCTG |
| 4.445257e-02 | GCNGCNC\|GNGCNGC |
| 4.402429e-02 | CNNGCNGC\|GCNGCNNG |
| 4.110281e-02 | CGNNCNG\|CNGNNCG |
| 4.090604e-02 | CTGGCNC\|GNGCCAG |

Supplementary Table 2: Heart active enhancer versus random k-mers

| Coefficient | k-mer |
|---|---|
| 4.092643e-02 | ATATNNAA\|TTNNATAT |
| 4.082406e-02 | ATATANA\|TNTATAT |
| 3.410699e-02 | AANATAT\|ATATNTT |
| 1.913669e-02 | TATNANAA\|TTNTNATA |
| 1.152353e-02 | ATATNTA\|TANATAT |
| 8.781651e-03 | ANATATA\|TATATNT |
| 8.651638e-03 | ATATTNT\|ANAATAT |
| 8.373656e-03 | ATATAA\|TTATAT |
| 6.013567e-03 | ANNATATA\|TATATNNT |
| 3.185595e-03 | ATATA\|TATAT |
| 2.389406e-03 | ATATANNT\|ANNTATAT |
| 5.172689e-04 | AANANTAT\|ATANTNTT |
| 1.364710e-04 | ATATNNAT\|ATNNATAT |
| 4.621480e-05 | CNTATA\|TATANG |
| 4.850692e-07 | AATNNTAT\|ATANNATT |

Supplementary Table 3: Kidney active enhancer versus random k-mers

| Coefficient | k-mer |
|---|---|
| 7.844098e-02 | ATTTNNAT\|ATNNAAAT |
| 6.227846e-02 | AAAATC\|GATTTT |
| 3.799466e-02 | AATATNNA\|TNNATATT |
| 1.514132e-02 | ATANAAT\|ATTNTAT |
| 1.126232e-02 | AANNATAT\|ATATNNTT |
| 9.860305e-03 | AATNNTAT\|ATANNATT |
| 4.661901e-03 | AANATAT\|ATATNTT |
| 3.743103e-03 | ATTNNATA\|TATNNAAT |
| 3.002073e-03 | ATATNAA\|TTNATAT |
| 6.330809e-04 | ATATTNNT\|ANNAATAT |

Supplementary Table 4: Liver active enhancer versus random k-mers

| Coefficient | k-mer |
|---|---|
| 2.287901e-02 | ATANNTAA\|TTANNTAT |
| 1.607827e-02 | ATATNTA\|TANATAT |
| 1.129433e-02 | ATATAA\|TTATAT |
| 9.325971e-03 | AATATNNA\|TNNATATT |
| 5.390299e-03 | ATANANAA\|TTNTNTAT |
| 2.456531e-03 | AANATNTA\|TANATNTT |
| 1.127811e-03 | ANATATNA\|TNATATNT |
| 1.026810e-03 | ATATA\|TATAT |
| 9.111476e-04 | ANNATATA\|TATATNNT |
| 6.708790e-04 | TATNANAA\|TTNTNATA |
| 1.751466e-07 | ANATATA\|TATATNT |

Supplementary Table 5: Limb active enhancer versus random k-mers

| Coefficient | k-mer |
|---|---|
| 5.287803e-02 | ATATANNT\|ANNTATAT |
| 3.219219e-02 | ATNNAATA\|TATTNNAT |
| 2.952235e-02 | AATNNTAT\|ATANNATT |
| 2.121272e-02 | ATATAA\|TTATAT |
| 1.992908e-02 | AANATAT\|ATATNTT |
| 1.321708e-02 | ATATNNAA\|TTNNATAT |
| 1.032477e-02 | ATATNNAT\|ATNNATAT |
| 7.143787e-03 | TATNANAA\|TTNTNATA |
| 3.685859e-03 | ATATTNT\|ANAATAT |
| 7.770563e-04 | ATNTATNA\|TNATANAT |
| 6.815450e-04 | ATAANAT\|ATNTTAT |
| 5.183059e-05 | CNTATA\|TATANG |

Supplementary Table 6: Lung active enhancer versus random k-mers

| Coefficient | k-mer |
|---|---|
| 7.705880e-02 | CNTATA\|TATANG |
| 3.225029e-02 | CATATA\|TATATG |
| 1.789592e-02 | ATATANA\|TNTATAT |
| 2.945417e-03 | GAGTTNNA\|TNNAACTC |
| 5.286091e-07 | ATATANNT\|ANNTATAT |

Supplementary Table 7: Forebrain active enhancer versus random k-mers

| Coefficient | k-mer |
|---|---|
| 4.967507e-02 | AAAGTT\|AACTTT |
| 3.015293e-02 | CNTATA\|TATANG |
| 2.849713e-02 | ATATANA\|TNTATAT |
| 2.261469e-02 | AAACTT\|AAGTTT |
| 1.191651e-02 | GAGTTNNA\|TNNAACTC |
| 8.910725e-03 | AACTTNT\|ANAAGTT |
| 8.839285e-03 | CATATA\|TATATG |

Supplementary Table 8: Midbrain active enhancer versus random k-mers

| Coefficient | k-mer |
|---|---|
| 5.341277e-02 | CNTATA\|TATANG |
| 3.697316e-02 | ANATATA\|TATATNT |
| 3.657011e-02 | ATATANA\|TNTATAT |
| 2.917223e-02 | AANATAT\|ATATNTT |
| 3.764890e-03 | AAAGTT\|AACTTT |
| 1.539902e-04 | AANANTAT\|ATANTNTT |
| 2.499315e-05 | AAAANTT\|AANTTTT |

Supplementary Table 9: Hindbrain active enhancer versus random k-mers

| Coefficient | k-mer |
|---|---|
| 1.882018e-01 | CTTATC\|GATAAG |
| 1.638524e-01 | AGGTCA\|TGACCT |
| 9.075613e-02 | AAAATAG\|CTATTTT |
| 8.853232e-02 | CAAGGNCA\|TGNCCTTG |
| 7.665507e-02 | AANATAG\|CTATNTT |
| 5.752587e-02 | TATNTNTA\|TANANATA |
| 5.717503e-02 | ANATAGC\|GCTATNT |
| 5.676097e-02 | TATTTNTA\|TANAAATA |
| 4.573678e-02 | CNTGTCA\|TGACANG |
| 4.471814e-02 | ANGTCAC\|GTGACNT |
| 4.285185e-02 | CNGATA\|TATCNG |
| 3.984399e-02 | ANNAATAG\|CTATTNNT |
| 3.829973e-02 | GNTGACA\|TGTCANC |
| 3.185298e-02 | CNTATC\|GATANG |
| 3.184707e-02 | GTCANCA\|TGNTGAC |
| 2.997837e-02 | AGATANNG\|CNNTATCT |
| 2.782536e-02 | CNNGTGAC\|GTCACNNG |
| 2.629641e-02 | ATTNNTAG\|CTANNAAT |
| 2.530363e-02 | ANGGTCA\|TGACCNT |
| 2.286028e-02 | AGTNCTG\|CAGNACT |

Supplementary Table 10: Heart active enhancer k-mers

| Coefficient | k-mer |
|---|---|
| 1.020531e-01 | CNNTGACC\|GGTCANNG |
| 9.948961e-02 | ATTAAC\|GTTAAT |
| 9.763349e-02 | AGGTCA\|TGACCT |
| 7.191567e-02 | ANNGGTCA\|TGACCNNT |
| 6.905743e-02 | GGGTCA\|TGACCC |
| 6.754502e-02 | CTCCCAC\|GTGGGAG |
| 6.525444e-02 | AAGTTNNA\|TNNAACTT |
| 6.500284e-02 | GATTTA\|TAAATC |
| 6.387318e-02 | CATNAANC\|GNTTNATG |
| 3.620010e-02 | GTNAATNA\|TNATTNAC |
| 3.502837e-02 | CTGACC\|GGTCAG |
| 3.428181e-02 | CCATNAA\|TTNATGG |
| 3.282915e-02 | AGTTNANG\|CNTNAACT |
| 3.228190e-02 | CNNGGTCA\|TGACCNNG |
| 3.212338e-02 | CAGGTNA\|TNACCTG |
| 3.144703e-02 | GGTTAA\|TTAACC |
| 2.510691e-02 | CCNCCCNC\|GNGGGNGG |
| 2.445600e-02 | CNTGACC\|GGTCANG |
| 2.394907e-02 | GTTAANNA\|TNNTTAAC |
| 2.385617e-02 | AAGTTNA\|TNAACTT |

Supplementary Table 11: Kidney active enhancer k-mers

| Coefficient | k-mer |
|---|---|
| 4.418726e-01 | CTTATC\|GATAAG |
| 3.842613e-01 | AGATAA\|TTATCT |
| 2.415544e-01 | TGATAA\|TTATCA |
| 2.166643e-01 | AGATANG\|CNTATCT |
| 1.408881e-01 | GGTGGNNC\|GNNCCACC |
| 1.024764e-01 | CTGATA\|TATCAG |
| 8.786420e-02 | ANAGATA\|TATCTNT |
| 7.655353e-02 | AGTTCNA\|TNGAACT |
| 7.446016e-02 | AGATANNG\|CNNTATCT |
| 6.929535e-02 | AGTTNNAG\|CTNNAACT |
| 5.718874e-02 | GAGTTCNA\|TNGAACTC |
| 5.654989e-02 | GAGATA\|TATCTC |
| 5.442684e-02 | CNACCAC\|GTGGTNG |
| 4.427998e-02 | AGATANNA\|TNNTATCT |
| 4.313449e-02 | CNTTATC\|GATAANG |
| 4.220056e-02 | AGATANC\|GNTATCT |
| 4.088724e-02 | CACACCC\|GGGTGTG |
| 4.069697e-02 | ANNAGATA\|TATCTNNT |
| 3.259381e-02 | ANGNCTAC\|GTAGNCNT |
| 2.730205e-02 | GTAGANNA\|TNNTCTAC |

Supplementary Table 12: Liver active enhancer k-mers

| Coefficient | k-mer |
|---|---|
| 8.756674e-02 | CTTGGC\|GCCAAG |
| 8.739274e-02 | ANTTTCCA\|TGGAAANT |
| 6.196472e-02 | GCCAANNC\|GNNTTGGC |
| 4.770059e-02 | CNTGGNCC\|GGNCCANG |
| 4.538072e-02 | ACNTNTGG\|CCANANGT |
| 3.698461e-02 | TGGAAAA\|TTTTCCA |
| 3.471553e-02 | ACATCTG\|CAGATGT |
| 3.331488e-02 | TGGAAANA\|TNTTTCCA |
| 3.251135e-02 | TTCCANA\|TNTGGAA |
| 2.754918e-02 | CATNTGG\|CCANATG |
| 2.203030e-02 | AAACTT\|AAGTTT |
| 2.069077e-02 | CCAGANNC\|GNNTCTGG |
| 1.509852e-02 | ANACCNCA\|TGNGGTNT |
| 1.388453e-02 | CNATAAA\|TTTATNG |
| 1.365222e-02 | ANNCCAGA\|TCTGGNNT |
| 9.218143e-03 | CCNATAA\|TTATNGG |
| 8.995484e-03 | CANACCC\|GGGTNTG |
| 7.366801e-03 | CNNGCCAA\|TTGGCNNG |
| 3.353828e-03 | GGCCCA\|TGGGCC |
| 3.290024e-03 | CANCTGG\|CCAGNTG |

Supplementary Table 13: Limb active enhancer k-mers

| Coefficient | k-mer |
|---|---|
| 1.406340e-01 | TAAACA\|TGTTTA |
| 8.770747e-02 | GTAAAC\|GTTTAC |
| 8.654490e-02 | TGTTTANA\|TNTAAACA |
| 7.529245e-02 | CAAACA\|TGTTTG |
| 5.069352e-02 | ANATTCC\|GGAATNT |
| 5.057327e-02 | ACTTCNNG\|CNNGAAGT |
| 4.514979e-02 | AAGTNCT\|AGNACTT |
| 3.844192e-02 | AAACTT\|AAGTTT |
| 3.057111e-02 | ANNTAAAC\|GTTTANNT |
| 2.795234e-02 | AGNAAACA\|TGTTTNCT |
| 2.753212e-02 | ATAAACA\|TGTTTAT |
| 2.531748e-02 | TTTGNTNA\|TNANCAAA |
| 2.358802e-02 | ACTTNTG\|CANAAGT |
| 2.319156e-02 | TTNCANAA\|TTNTGNAA |
| 2.287478e-02 | ANNGTTTA\|TAAACNNT |
| 2.190120e-02 | GTTTGNNC\|GNNCAAAC |
| 2.184071e-02 | AAGTNNCT\|AGNNACTT |
| 2.159117e-02 | ANTNTTTG\|CAAANANT |
| 2.139472e-02 | ACATTCC\|GGAATGT |
| 2.057346e-02 | ACTTCNT\|ANGAAGT |

Supplementary Table 14: Lung active enhancer k-mers

| Coefficient | k-mer |
|---|---|
| 3.284869e-01 | CAGATGG\|CCATCTG |
| 1.747920e-01 | CAATTA\|TAATTG |
| 1.558061e-01 | ATTNGCA\|TGCNAAT |
| 1.048507e-01 | CNAATTA\|TAATTNG |
| 8.073584e-02 | TAATTA\|TAATTA |
| 7.885150e-02 | AATTAG\|CTAATT |
| 6.729461e-02 | ATGNNAAT\|ATTNNCAT |
| 5.550338e-02 | ACANATNG\|CNATNTGT |
| 4.861347e-02 | ATGCNAA\|TTNGCAT |
| 4.517273e-02 | ATGAAT\|ATTCAT |
| 4.471312e-02 | ANNATATG\|CATATNNT |
| 3.985233e-02 | TGCCAA\|TTGGCA |
| 3.959699e-02 | CATTTNNA\|TNNAAATG |
| 3.744615e-02 | ANNTGGCA\|TGCCANNT |
| 3.647300e-02 | AATTANNC\|GNNTAATT |
| 3.586182e-02 | CATATG\|CATATG |
| 3.507460e-02 | ATTTNNAT\|ATNNAAAT |
| 3.461155e-02 | AATTANC\|GNTAATT |
| 3.446622e-02 | CTNATTA\|TAATNAG |
| 2.912524e-02 | AATNNGCA\|TGCNNATT |

Supplementary Table 15: Forebrain active enhancer k-mers

| Coefficient | k-mer |
|---|---|
| 1.194402e-01 | TATTCA\|TGAATA |
| 1.092740e-01 | AATAAT\|ATTATT |
| 8.937580e-02 | ATGNNAAT\|ATTNNCAT |
| 8.665814e-02 | GATNAAA\|TTTNATC |
| 8.247386e-02 | ATTTNNAT\|ATNNAAAT |
| 6.004712e-02 | ATTTNAA\|TTNAAAT |
| 4.700935e-02 | ATCAAA\|TTTGAT |
| 3.942919e-02 | ATTATNC\|GNATAAT |
| 3.730211e-02 | ATTATNNA\|TNNATAAT |
| 2.954801e-02 | AATNNAAT\|ATTNNATT |
| 2.938446e-02 | ATNAAANG\|CNTTTNAT |
| 2.914561e-02 | AAAATC\|GATTTT |
| 2.613495e-02 | CTNATTA\|TAATNAG |
| 2.508567e-02 | ATNAATA\|TATTNAT |
| 2.357498e-02 | AATCNNTT\|AANNGATT |
| 2.307671e-02 | ATNAAAG\|CTTTNAT |
| 2.175419e-02 | ATNNATTA\|TAATNNAT |
| 1.905055e-02 | TAATGA\|TCATTA |
| 1.504929e-02 | ATNAANAG\|CTNTTNAT |
| 1.206887e-02 | ATTANNNC\|GNNNTAAT |

Supplementary Table 16: Midbrain active enhancer k-mers

| Coefficient | k-mer |
|---|---|
| 6.310558e-02 | TAATNAA\|TTNATTA |
| 4.995234e-02 | ATNNATCA\|TGATNNAT |
| 4.090126e-02 | CNNTGGCA\|TGCCANNG |
| 3.064466e-02 | CGNNNAGC\|GCTNNNCG |
| 3.004314e-02 | AGCNNNCG\|CGNNNGCT |
| 2.948802e-02 | AATTGNNT\|ANNCAATT |
| 2.872896e-02 | CNCGCA\|TGCGNG |
| 2.674954e-02 | CTAATNA\|TNATTAG |
| 2.322091e-02 | CANCGC\|GCGNTG |
| 2.315027e-02 | ATNCATC\|GATGNAT |
| 1.798161e-02 | GCATCNNC\|GNNGATGC |
| 1.349962e-02 | CNCTGCNG\|CNGCAGNG |
| 1.281059e-02 | ATGGCA\|TGCCAT |
| 1.233782e-02 | CGNTGC\|GCANCG |
| 1.200482e-02 | GCNGCTNC\|GNAGCNGC |
| 1.046248e-02 | ANGATGC\|GCATCNT |
| 1.003060e-02 | CTGCNGNC\|GNCNGCAG |
| 9.450373e-03 | CATNNATC\|GATNNATG |
| 8.122928e-03 | GCAGNGC\|GCNCTGC |
| 7.055018e-03 | AGCGA\|TCGCT |

Supplementary Table 17: Hindbrain active enhancer k-mers

| Coefficient | k-mer |
|---|---|
| 7.108817e-02 | AAATAG\|CTATTT |
| 6.639535e-02 | GAAAANNA\|TNNTTTTC |
| 4.611863e-02 | ATTTNNAG\|CTNNAAAT |
| 1.874686e-02 | TAANAANA\|TNTTNTTA |
| 1.087803e-02 | ANAAATA\|TATTTNT |
| 7.252695e-03 | AAAATA\|TATTTT |
| 5.869543e-03 | TAAAANNA\|TNNTTTTA |
| 4.795092e-03 | AAANAGNA\|TNCTNTTT |
| 4.711524e-03 | ANNTTTAG\|CTAAANNT |
| 3.249960e-03 | TATTTNNA\|TNNAAATA |
| 2.100815e-03 | TTTCNAA\|TTNGAAA |

Supplementary Table 18: Heart active/primed enhancer k-mers

| Coefficient | k-mer |
|---|---|
| 2.595118e-02 | AGGAANT\|ANTTCCT |
| 2.510978e-02 | CANGAAA\|TTTCNTG |
| 2.186890e-02 | CANNAAGC\|GCTTNNTG |
| 1.809457e-02 | ANATTCC\|GGAATNT |
| 1.584056e-02 | GNAAACA\|TGTTTNC |
| 1.066339e-02 | AANTTCC\|GGAANTT |
| 7.775262e-03 | ATTTCC\|GGAAAT |
| 7.707218e-03 | AGGNAGGA\|TCCTNCCT |
| 3.535195e-03 | CNGGAAA\|TTTCCNG |
| 2.838993e-03 | GNAGGAA\|TTCCTNC |
| 1.873522e-03 | CAGGAAG\|CTTCCTG |
| 1.480342e-03 | AAANNAAC\|GTTNNTTT |
| 6.165058e-04 | AAACAA\|TTGTTT |
| 4.955818e-04 | GCNGGAA\|TTCCNGC |

Supplementary Table 19: Kidney active/primed enhancer k-mers

| Coefficient | k-mer |
|---|---|
| 2.726096e-02 | CNCAGNAA\|TTNCTGNG |
| 2.466940e-02 | ANACAAG\|CTTGTNT |
| 2.289408e-02 | GCCTNAC\|GTNAGGC |
| 1.448976e-02 | ACANGCC\|GGCNTGT |
| 1.275854e-02 | AGGCCNNT\|ANNGGCCT |
| 7.445750e-03 | GGNCNACA\|TGTNGNCC |
| 4.608107e-03 | AGCTNAG\|CTNAGCT |
| 1.362636e-03 | ANAGNAAC\|GTTNCTNT |

Supplementary Table 20: Liver active/primed enhancer k-mers

| Coefficient | k-mer |
|---|---|
| 4.946880e-02 | AGNAAANC\|GNTTTNCT |
| 4.142856e-02 | AAANAGNC\|GNCTNTTT |
| 3.357433e-02 | TGGAAA\|TTTCCA |
| 3.274859e-02 | ANNTTTCC\|GGAAANNT |
| 2.919853e-02 | ANNTAAAA\|TTTTANNT |
| 2.623187e-02 | ANTTTCC\|GGAAANT |
| 1.958511e-02 | GAANGNAA\|TTNCNTTC |
| 1.947151e-02 | AGTTNNTG\|CANNAACT |
| 1.596530e-02 | GNTTTNCA\|TGNAAANC |
| 1.355715e-02 | TGAANNAA\|TTNNTTCA |
| 1.086766e-02 | TTNAGNAA\|TTNCTNAA |
| 1.032791e-02 | AGANTTNC\|GNAANTCT |
| 9.988572e-03 | AAANCAA\|TTGNTTT |
| 9.082434e-03 | TGNNGAAA\|TTTCNNCA |
| 7.522810e-03 | AANCTCT\|AGAGNTT |
| 5.293159e-03 | AAANNTCT\|AGANNTTT |
| 3.609649e-03 | AGNAAANA\|TNTTTNCT |
| 3.455022e-03 | ANAGGAA\|TTCCTNT |
| 2.498589e-03 | ATTTNCNT\|ANGNAAAT |
| 6.208859e-04 | GGAAANA\|TNTTTCC |

Supplementary Table 21: Limb active/primed enhancer k-mers

| Coefficient | k-mer |
|---|---|
| 3.954595e-02 | GNAAACA\|TGTTTNC |
| 3.450363e-02 | AGNAAAA\|TTTTNCT |
| 3.376044e-02 | AGGAANT\|ANTTCCT |
| 2.923563e-02 | AGNAAAT\|ATTTNCT |
| 1.775037e-02 | ANAGGAA\|TTCCTNT |
| 1.648291e-02 | AGANNAAA\|TTTNNTCT |
| 1.585723e-02 | AAACANT\|ANTGTTT |
| 1.585384e-02 | ATTTCC\|GGAAAT |
| 1.367495e-02 | AAANAGA\|TCTNTTT |
| 1.222750e-02 | CAGGAAG\|CTTCCTG |
| 1.202030e-02 | AANNAAAG\|CTTTNNTT |
| 1.124490e-02 | AAACANNC\|GNNTGTTT |
| 1.064884e-02 | AANAAANG\|CNTTTNTT |
| 6.117277e-03 | ANTTTCC\|GGAAANT |
| 5.464630e-03 | AGNAAAC\|GTTTNCT |
| 5.254366e-03 | TTCNAAA\|TTTNGAA |
| 3.708424e-03 | TGTTTNA\|TNAAACA |
| 3.179645e-03 | AAAGNAA\|TTNCTTT |
| 2.036569e-03 | AAGAAA\|TTTCTT |
| 1.867751e-03 | TNTAAANA\|TNTTTANA |

Supplementary Table 22: Lung active/primed enhancer k-mers

| Coefficient | k-mer |
|---|---|
| 1.590400e-01 | CAGATGG\|CCATCTG |
| 1.970634e-02 | ANAAAGC\|GCTTTNT |
| 1.873085e-02 | ACANANGG\|CCNTNTGT |
| 1.862789e-02 | CANANAGC\|GCTNTNTG |
| 1.855113e-02 | AANAAAGG\|CCTTTNTT |
| 1.467011e-02 | ACAAANG\|CNTTTGT |
| 1.132120e-02 | ACAAAG\|CTTTGT |
| 8.978300e-03 | CNGCAGC\|GCTGCNG |
| 7.756535e-03 | ANGCAGC\|GCTGCNT |
| 7.199301e-03 | AGNCAGC\|GCTGNCT |
| 7.149984e-03 | AGAGCT\|AGCTCT |
| 7.121278e-03 | ACAAAGG\|CCTTTGT |
| 5.402761e-03 | CANATGG\|CCATNTG |
| 4.022826e-03 | GANNGAAA\|TTTCNNTC |
| 2.913987e-03 | AANANAGA\|TCTNTNTT |
| 2.079053e-03 | AGCTGNNA\|TNNCAGCT |
| 1.919643e-03 | GNGAGANA\|TNTCTCNC |
| 1.884114e-03 | GCTGNCA\|TGNCAGC |
| 1.654904e-03 | AANGAGA\|TCTCNTT |
| 1.433402e-03 | ANANAGCC\|GGCTNTNT |

Supplementary Table 23: Forebrain active/primed enhancer k-mers

| Coefficient | k-mer |
|---|---|
| 2.916380e-02 | GAAANGC\|GCNTTTC |
| 1.987357e-02 | CNGCAGC\|GCTGCNG |
| 1.884978e-02 | ATTNGCA\|TGCNAAT |
| 1.586346e-02 | GCAGCC\|GGCTGC |
| 1.317559e-02 | ANAAANGC\|GCNTTTNT |
| 1.033002e-02 | GTTGCNA\|TNGCAAC |
| 7.934415e-03 | CNGCNGC\|GCNGCNG |
| 6.761922e-03 | AGCNGCNG\|CNGCNGCT |
| 4.859020e-03 | AGCTGC\|GCAGCT |
| 3.454828e-03 | ANNGGAAA\|TTTCCNNT |
| 3.396658e-03 | CNGNCAGC\|GCTGNCNG |
| 2.929552e-03 | AAANAGC\|GCTNTTT |
| 2.127981e-03 | AAANGAG\|CTCNTTT |
| 1.505571e-03 | GCNGANAA\|TTNTCNGC |
| 1.428004e-03 | ATTTGC\|GCAAAT |

Supplementary Table 24: Midbrain active/primed enhancer k-mers

| Coefficient | k-mer |
|---|---|
| 8.847729e-02 | ACAAAG\|CTTTGT |
| 6.154593e-02 | ACAAANG\|CNTTTGT |
| 2.916447e-02 | GNAAATNA\|TNATTTNC |
| 2.555737e-02 | AAANNGCT\|AGCNNTTT |
| 2.532082e-02 | ATTTNCA\|TGNAAAT |
| 2.463892e-02 | ANAAANGC\|GCNTTTNT |
| 2.092278e-02 | GCTGCA\|TGCAGC |
| 2.031603e-02 | AGCNGCNG\|CNGCNGCT |
| 1.533200e-02 | GAANAAA\|TTTNTTC |
| 1.471883e-02 | CNGCAGC\|GCTGCNG |
| 1.417746e-02 | GAAANGC\|GCNTTTC |
| 1.376909e-02 | ATTTCNNT\|ANNGAAAT |
| 1.224286e-02 | CNTTTNTC\|GANAAANG |
| 1.195203e-02 | AAANAGC\|GCTNTTT |
| 5.577006e-03 | AAGCNGC\|GCNGCTT |
| 4.821936e-03 | ATTTCC\|GGAAAT |
| 4.380671e-03 | AANGANGC\|GCNTCNTT |
| 4.255284e-03 | ANGCAGC\|GCTGCNT |
| 3.652331e-03 | AACAANG\|CNTTGTT |
| 3.599566e-03 | GAAANAAA\|TTTNTTTC |

Supplementary Table 25: Hindbrain active/primed enhancer k-mers

| | ATAC | H3K27ac | H3K4me1 | H3K4me3 | H3K9me3 | H3K27me3 | RNA | Control |
|---|---|---|---|---|---|---|---|---|
| Forebrain | ENCFF695FLH<br>ENCFF572CMB | ENCFF761HMI<br>ENCFF236VKK | ENCFF924YFS<br>ENCFF551NRH | ENCFF911RYA<br>ENCFF755BPV | ENCFF204RVT<br>ENCFF403OKB | ENCFF460TKS<br>ENCFF926ANX | ENCFF798GDC<br>ENCFF598ESF | ENCFF901RYX<br>ENCFF880SQD |
| Hindbrain | | ENCFF413NTF<br>ENCFF156DKT | ENCFF989KUF<br>ENCFF891VNO | ENCFF884HYZ<br>ENCFF955BLP | ENCFF170JCI<br>ENCFF485DDP | ENCFF642MWW<br>ENCFF187EVU | ENCFF630JFR<br>ENCFF672XLI | ENCFF123XKF<br>ENCFF801CNP |
| Midbrain | ENCFF060YWL<br>ENCFF100IBX | ENCFF054UNT<br>ENCFF892TDG | ENCFF526EGV<br>ENCFF190ZOW | ENCFF960GUP<br>ENCFF196TGU | ENCFF986SRM<br>ENCFF958RLB | ENCFF318OLQ<br>ENCFF626OTI | ENCFF785FFD<br>ENCFF056VIQ | ENCFF643WUI<br>ENCFF916TFR |
| Heart | | ENCFF576RBC<br>ENCFF261KJB | ENCFF839EWF<br>ENCFF156UJE | ENCFF978DEK<br>ENCFF292UUU | ENCFF502UOP<br>ENCFF262FYW | ENCFF275AKZ<br>ENCFF420BPA | ENCFF951VYO<br>ENCFF143ATC | ENCFF487AIV<br>ENCFF642FOT |
| Kidney | ENCFF036RNM<br>ENCFF477SZP | ENCFF650CDV<br>ENCFF831JKI | ENCFF071PID<br>ENCFF167AOI | ENCFF262SUZ<br>ENCFF407ALD | ENCFF148BAU<br>ENCFF110BCP | ENCFF139UWI<br>ENCFF977IBX | ENCFF258QXM<br>ENCFF522LYF | ENCFF627RIT<br>ENCFF285XPR |
| Limb | | ENCFF156ZFI<br>ENCFF052AZF | ENCFF173RDF<br>ENCFF295BOL | ENCFF148HFT<br>ENCFF536ZMQ | ENCFF894UZN<br>ENCFF011YLZ | ENCFF003PZM<br>ENCFF518UCO | ENCFF445SGU<br>ENCFF223JGE | ENCFF031OYA<br>ENCFF514GVM |
| Liver | | ENCFF027UTM<br>ENCFF826GUO | ENCFF783RDN<br>ENCFF141HFI | ENCFF031RFH<br>ENCFF492LOP | ENCFF477WQY<br>ENCFF858QZZ | ENCFF869LGO<br>ENCFF031MFD | ENCFF789CGL<br>ENCFF043NJH | ENCFF209HPI<br>ENCFF634NAV |
| Lung | | ENCFF561MSI<br>ENCFF497ZYH | ENCFF338GKD<br>ENCFF555LDI | ENCFF401RKM<br>ENCFF669NCH | ENCFF380FFY<br>ENCFF222SXN | ENCFF788XZB<br>ENCFF801BAX | ENCFF870FAN<br>ENCFF699ZJN | ENCFF079JAD<br>ENCFF269JKO |

Supplementary Table 26: ENCODE bam accession codes

| | | | | |
|---|---|---|---|---|
| Hindbrain | ENCFF761JHA | ENCFF181OVJ | ENCFF213OUA | ENCFF709CLT |
| Heart | ENCFF829XFO | ENCFF694SPD | ENCFF385WYM | ENCFF051GLX |
| Limb | ENCFF629FRA | ENCFF222KJL | ENCFF488EFS | ENCFF917DTZ |
| Liver | ENCFF329VCX | ENCFF290ZBP | ENCFF341HRL | ENCFF489XAT |
| Lung | ENCFF507MHY | ENCFF217WJG | ENCFF159YPF | ENCFF940GZL |

Supplementary Table 27: ENCODE ATAC fastq accession codes
